# Dissecting endocytic mechanisms reveals new molecular targets to enhance sodium iodide symporter activity with clinical relevance to radioiodide therapy

**DOI:** 10.1101/2023.05.22.541733

**Authors:** Martin L. Read, Katie Brookes, Ling Zha, Selvambigai Manivannan, Jana Kim, Merve Kocbiyik, Alice Fletcher, Caroline M. Gorvin, George Firth, Gilbert O. Fruhwirth, Juan P. Nicola, Sissy Jhiang, Matthew D. Ringel, Moray J. Campbell, Kavitha Sunassee, Philip J. Blower, Kristien Boelaert, Hannah R. Nieto, Vicki E. Smith, Christopher J. McCabe

## Abstract

The sodium/iodide symporter (NIS) frequently shows diminished plasma membrane (PM) targeting in differentiated thyroid cancer (DTC), resulting in suboptimal radioiodide (RAI) treatment and poor prognosis. The mechanisms which govern the endocytosis of NIS away from the PM are ill-defined. Here, we challenged the hypothesis that new mechanistic understanding of NIS endocytosis would facilitate prediction of patient outcomes and enable specific drug modulation of RAI uptake in vivo. Through mutagenesis, NanoBiT interaction assays, cell surface biotinylation assays, RAI uptake and NanoBRET, we identify an acidic dipeptide within the NIS C-terminus which mediates binding to the σ2 subunit of the Adaptor Protein 2 (AP2) heterotetramer. We discovered that the FDA-approved drug chloroquine modulates NIS accumulation at the PM in a functional manner that is AP2 dependent. In vivo, chloroquine treatment of BALB/c mice significantly enhanced thyroidal uptake of ^99m^Tc pertechnetate in combination with the histone deacetylase (HDAC) inhibitor SAHA, accompanied by increased thyroidal NIS mRNA. Bioinformatic analyses validated the clinical relevance of AP2 genes with disease-free survival in RAI-treated DTC, enabling construction of an AP2 gene-related risk score classifier for predicting recurrence. We propose that NIS internalisation is orchestrated by the interaction of a C-terminal diacidic motif with AP2σ2, together with the proto-oncogene PBF acting via AP2μ2. Given that NIS internalisation was specifically druggable in vivo, our data provide new translatable potential for improving RAI therapy using FDA-approved drugs in patients with aggressive thyroid cancer.

**Summary:** We delineate the role of endocytic genes in regulating NIS activity at the plasma membrane and highlight the potential for systemic targeting of endocytosis to enhance radioiodine effectiveness in radioiodine-refractory cancer cells.

## INTRODUCTION

β-emitting radioiodide (^131^I) has been utilised for over 80 years to safely, efficiently, and specifically destroy remaining thyroid cancer cells post-surgery and to target metastases (1). Patients with radioiodide-resistant (RAIR) thyroid cancer, particularly those with metastatic disease, have a life expectancy of only 3–5 years and represent a group for whom there is a clear unmet medical need (2). The sodium/ iodide symporter (NIS) is the sole transporter of iodide into human cells; tumoural radioiodide uptake is diminished in 25-50% of thyroid cancer patients, due to reduced expression and mislocalisation away from the plasma membrane (PM) (3–5), its only site of transport activity. Whilst drugs have been developed which restore NIS mRNA and protein expression in preclinical models (6) and in subsets of patients (7), understanding and manipulating firstly how the NIS protein is trafficked to the PM, and secondly how it is internalised away from it, remains essential in enhancing the function of NIS in radioiodide treatment. The endocytosis of membrane transporters and symporters is a key determinant of actual transport activity, and yet for NIS, the mechanism(s) governing its endocytosis are largely unknown.

Importantly, knowledge of NIS trafficking and endocytosis could also feed into other clinical settings, particularly breast cancer. NIS expression is inappropriately activated in ∼60 to 80% of breast tumours, including triple-negative breast cancers (TNBC) and brain metastases (5,8,9). Although radioiodide uptake into breast tumours and metastases has been demonstrated, levels of uptake are insufficient to achieve a therapeutic effect (10), as NIS is generally found in a non-functional intracellular location (5,9,10). We have recently identified a clear pathway via which we can drive NIS to the PM *in vitro* (11). However, there is no cogent understanding of the mechanisms which control the endocytosis of NIS at present (12).

More generally, the processes that govern the relationship between membrane transport of a substrate and when the membrane transporter itself is actually internalised are incompletely understood. Our group previously identified the proto-oncogene PTTG1 Binding Factor (PBF) to be a NIS-interacting protein, and to be capable of inducing NIS endocytosis when over- expressed (13–15). Both PBF and NIS have long C-terminal ‘tails’ which are able to bind other proteins. PBF is over-expressed in thyroid and breast cancer (16,17), and cellular expression results in decreased NIS localisation at the PM and reduced radioiodide uptake (13–15). PBF has a canonical YXXΦ tyrosine-based endocytosis motif at its C-terminus (174YARF177). Abrogating PBF Y174 (Y174A mutant) prevents NIS binding, consequently increasing NIS localisation at the PM and enhancing radioiodide uptake (15). PBF Y174 is also phosphorylated, and depletion of PBF pY174 using Src kinase inhibitors similarly restores radioiodide uptake, suggesting that PBF phosphorylation status mediates its regulation of NIS function (15).

The basolateral targeting of NIS has been investigated in several studies, and motifs responsible for interacting with the AP1 machinery defined (18–20). Moreover, NIS sorting to and retention at the PM requires additional motifs (21,22). However, it is not currently known how NIS endocytoses away from the PM after its trafficking there. Herein, we sought to define the mechanisms of NIS endocytosis in thyroid cancer cells, with the hypothesis that transiently inhibiting the movement of NIS away from the PM would result in significantly enhanced cellular radionuclide uptake. We identify that a diacidic/dileucine motif in the C-terminus of NIS governs its ability to interact with the σ2 subunit of AP2 and that AP2 modulates the interaction between NIS and PBF. A detailed bioinformatic analysis further demonstrated extensive dysregulation of endocytosis genes in DTC, as well as enabling construction of a AP2 gene-related independent predictive risk model for recurrent thyroid cancer. Our study reveals that the FDA-approved drug chloroquine retains NIS at the PM. Critically, in BALB/c mice, a combination of CQ with the histone deacetylase (HDAC) inhibitor SAHA enhances thyroidal uptake of the radionuclide ^99m^Tc, suggesting that NIS internalisation may now be druggable in vivo.

## MATERIALS AND METHODS

### Cell culture

Thyroid (TPC-1, 8505C, SW1736) cancer cell lines were maintained in RPMI-1640 (Life Technologies), while HeLa and HEK293 cancer cells were maintained in DMEM (Sigma-Aldrich). Media was supplemented with 10% fetal bovine serum (FBS), penicillin (10^5^ U/l), and streptomycin (100 mg/l) and cell lines were maintained at 37°C and 5% CO2 in a humidified environment. Cell lines were obtained from ECACC (HEK293, HeLa) and DSMZ (8505C), while TPC-1 and SW1736 cell lines were kindly provided by Dr Rebecca Schweppe (University of Colorado). Cells were cultured at low passage, authenticated by short tandem repeat analysis (NorthGene; Supp Fig. S1) and tested for mycoplasma contamination (EZ-PCR kit; Geneflow). Stable TPC-1-NIS and 8505C-NIS cells were generated by transfection of parental TPC-1 or 8505C cells with pcDNA3.1-NIS. Geneticin-resistant monoclonal colonies were expanded following FACS single cell sorting (University of Birmingham Flow Cytometry Facility), and Western blotting used to confirm NIS expression.

### Human thyroid tissue

Human thyroid tissue was obtained with local ethics committee approval from the Human Biomaterials Resource Centre (University of Birmingham) and informed patient consent. Primary thyrocytes were isolated and cultured as described (23).

### Inhibitors and drugs

Chloroquine diphosphate (Sigma-Aldrich) was resuspended in PBS without calcium/magnesium (ThermoFisher). SAHA (Stratech Scientific) and Dynasore (Sigma-Aldrich) were resuspended in dimethyl sulfoxide (DMSO; Sigma-Aldrich). All drugs were diluted in RPMI-1640 medium (1:100; Life Technologies) prior to treatment of cells. For intraperitoneal administration (IP) in mice, SAHA was formulated in 5% DMSO, 40% PEG400, 5% Tween-80 and PBS.

### Nucleic acids and transfection

Plasmid containing human NIS cDNA with a HA-tag has been described (14). The QuikChange Site-directed Mutagenesis Kit (Agilent Technologies) was used to generate two NIS mutants [(L562A/L563A) and (E578A/E579A)], as well as two mutants of AP2S1 [(V88D) and (L103D)]. To construct plasmids for NanoBiT detection, AP2S1 and PBF cDNA were cloned into pcDNA3.1 containing LgBiT or amplified with the SmBiT tag prior to cloning into pcDNA3.1. NIS-NanoLuc (Nluc) cDNA was synthesized and subcloned into pcDNA3.1 by GeneArt (ThermoFisher Scientific). Professor Nevin Lambert (Georgia Regents University) kindly provided the NanoBRET PM marker Kras-Venus, as well as the Venus-tagged subcellular compartment markers Rab5, Rab7 and Rab11. Venus-tagged markers Rab1, Rab4, Rab6 and Rab8 were kindly provided by Professor Kevin Pfleger (University of Western Australia) (24). Further details on NanoBiT and NanoBRET plasmids are given in Supplementary Information. Plasmid DNA and siRNA transfections were performed with TransIT-LT1 (Mirus Bio) and Lipofectamine RNAiMAX (ThermoFisher Scientific) following standard protocols in accordance with the manufacturer’s guidelines.

### Western blotting, cell-surface biotinylation and RAI uptake

Western blotting, cell surface biotinylation assays and RAI (^125^I) uptake assays were performed as described previously (14,23). Blots were probed with specific antibodies against NIS (1:1000; Proteintech), AP2α1 (1:400; Antibodies.com), AP2μ2 (1:500; Novus Biologicals), HA (1:1000; BioLegend), Na,K-ATPase (1:500; Cell Signaling Technology); PICALM (1:1000; Cell Signaling Technology) and β-actin (1:2000; Sigma-Aldrich). HRP-conjugated secondary antibodies (Agilent Technologies) against either mouse or rabbit IgG were used at 1:10000 dilution.

### qPCR

Total RNA was extracted using the RNeasy Micro Kit (Qiagen) and reverse transcribed using the Reverse Transcription System (Promega). Mouse thyroid tissue was homogenized in buffer RLT using TissueLyser II (Qiagen; 2x 2 min cycles at 30 Hz) and 5 mm stainless steel beads. Expression of specific mRNAs was determined using 7500 Real-time PCR system (Applied Biosystems) as described previously (23). TaqMan qPCR assays used are listed in Supplementary Information.

### Immunofluorescence

24 hours post transfection, cells were washed with PBS and fixed for 15 minutes at RT in 4% paraformaldehyde/PBS. After rinsing in PBS and 0.1M glycine/PBS, cells were permeabilized in 0.1% saponin buffer. Incubation with a mixture of primary antibodies [mouse-anti-HA (1:100) and rabbit-anti-NIS (1:100)] was performed at RT for 1 hour. Cells were rinsed three times with saponin buffer before an 1 hour incubation with secondary antibodies (Alexa- Fluor-555-conjugated goat anti-rabbit or Alexa-Fluor-488-conjuated goat anti-mouse). Finally, cells were rinsed with saponin buffer (3x) and PBS (1x) and mounted onto slides using Prolong Gold anti-fade reagent with DAPI (Molecular Probes). Cells were viewed and images captured using 100X objective on a LSM 880 Airyscan confocal microscope. Images were analysed using FIJI software.

### NanoBiT and NanoBRET live cell assays

Cells were seeded in 6-well plates at a density of 3.5 x 10^5^ cells per well and transfected with 500 ng – 1 µg plasmid DNA. 24 hours post-transfection, cells were harvested and reseeded into white 96-well plates in phenol-red-free DMEM (Life Technologies). Furimazine (Promega) was added to each well in accordance with the manufacturer’s guidelines and readings taken at 120 second intervals for up to 40 minutes (PHERAstar FS microplate reader; BMG Labtech). In some experiments, cells were treated with CQ (8 hours) or DYN (24 hours) prior to addition of furimazine. The NanoBRET signal was calculated using standard protocols by dividing the acceptor emission at 618 nm by the donor emission at 460 nm.

### In vivo biodistribution

All animal experiments were performed in accordance with the Animals (Scientific Procedures) Act, 1986 with protocols approved by the Animal Welfare and Ethical Review Body for King’s College London (St Thomas’ Campus). Male BALB/c mice (8-10 weeks of age, *n* = 4 -18 animals/group, Charles River Laboratories) received either vehicle (PBS/DMSO), CQ (40-60 mg/kg/day), SAHA (100 mg/kg/day) or SAHA+CQ by IP injection for 4 consecutive days. CQ was administered 4 hours after SAHA. On day 4, mice were anaesthetized by isoflurane inhalation (3%, Animalcare, York, in O2) and maintained under isoflurane anesthesia during IV administration of ^99m^Tc-pertechnetate (0.5 MBq). After 30 minutes, mice were culled by anesthetic overdose and tissues harvested. Thyroid glands were removed using a dissecting microscope. Radioactivity was measured by gamma counting (1282 Compugamma; LKB Wallac).

### Differential gene expression and clinical data

Gene expression data and clinical information for papillary thyroid cancer (PTC) were downloaded from TCGA via cBioPortal (cbioportal.org/), FireBrowse (firebrowse.org) and NCI Genomic Data Commons (GDC; portal.gdc.cancer.gov/) (25–27). Bioinformatic approaches for thyroid TCGA and GEO data analyses are outlined (Supplementary Information).

### Statistical analyses

Statistical analyses were performed using IBM SPSS Statistics (Version 29), GraphPad Prism (Version 9.5) and Microsoft Excel. See Supplementary Information for details.

## RESULTS

### Endocytic factor AP2 is a critical regulator of functional NIS

To identify proteins which might bind NIS and influence its endocytosis we interrogated data from 2 mass spectrometry investigations (11,28) and compared this analysis with endocytosis pathway gene sets (29) and PathCards (30) (**Fig. 1A**; Supp Table S1). This allowed us to identify 14 potential NIS interactors associated with endocytotic pathways. Stratification of papillary thyroid cancer (THCA) TCGA expression data using quartile values (Q3Q4 vs Q1Q2) further highlighted the potential clinical relevance for 5 of these NIS interactors (AP2A2, ARF6, CTTN, HLA-A and RAB5C) on disease recurrence following RAI treatment (*P* < 0.05; **Fig. 1B**; Supp Fig. S2A and S2B). Based on this we selected the heterotetrameric AP2 adaptor complex with an established role in clathrin-mediated endocytosis and undertook an siRNA screen to investigate the role of AP2 subunits and AP2 associated kinase 1 (AAK1) on NIS function. Importantly, abrogating AP2 subunits α1 and μ2, as well as AAK1, significantly enhanced RAI uptake in NIS-stably expressing thyroid cancer cell lines TPC-1 (TPC-1-NIS) and 8505C (8505C-NIS) (**Fig. 1C**; Supp Fig. S2C), although depleting AP2σ2 had no effect. Abrogation of AP2 subunits in human primary thyrocytes had a similar effect in significantly enhancing RAI uptake (**Fig. 1C**), although AP2σ2 siRNA was associated with increased uptake. Interestingly, AP2α1 ablation was associated with increased NIS protein expression in TPC1-NIS and 8505C-NIS cells (**Fig. 1D**). Control experiments showed significant knockdown with siRNA targeting AP2 subunits and AAK1 (**Fig. 1E**), but negligible impact on NIS mRNA expression (Supp Fig. S2D).

**Figure 1.**
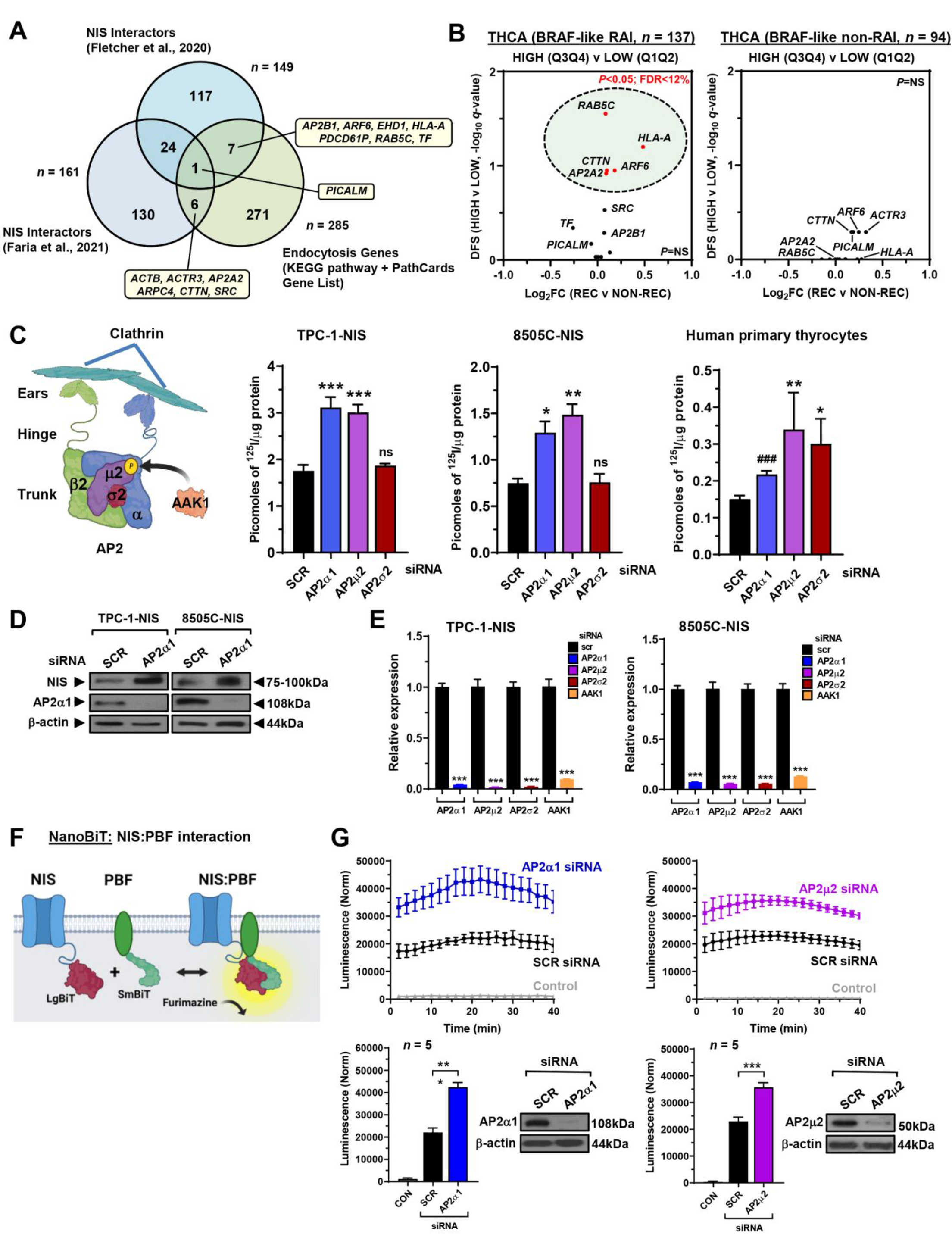
Modulation of AP2 expression increases RAI uptake. **A,** Venn diagram showing overlap in NIS interactors from 2 mass spectrometry investigations compared to known endocytosis genes (KEGG pathway + Pathcards). 14 top candidates are highlighted. **B,** Volcano plot illustrating log2FC [recurrent (REC) versus non-recurrent (NON-REC)] compared to *q*-value (-log base 10) for DFS in RAI-treated (*left*) and non-RAI treated (*right*) BRAF-like THCA cohorts for 14 endocytic genes [high (Q3Q4) versus low (Q1Q2) tumoral expression]. *P* < 0.05 (green circle). **C,** Schematic (*left*) depicting AP2 subunits and interaction with clathrin/ AAK1. Created with BioRender.com. (*right*) RAI uptake of TPC- 1-NIS cells, 8505C-NIS cells and human primary thyrocytes transfected with siRNA specific for indicated AP2 genes. **D,** Western blot analysis of NIS and AP2α1 protein in TPC-1-NIS and 8505C- NIS cells transfected with AP2α1 siRNA. **E,** Relative mRNA levels of AP2 genes and AAK1 in TPC- 1-NIS and 8505C-NIS cells transfected with siRNA specific for indicated AP2 genes and AAK1. **F,** Schematic depicting NanoBiT assay to detect protein: protein interaction between NIS tagged with LgBiT and PBF tagged with SmBiT. The NanoLuc luciferase enzyme (LgBiT + SmBiT) relies on the substrate furimazine to produce high intensity, glow-type luminescence. **G,** NanoBiT evaluation (*upper*) of protein: protein interaction between NIS and PBF in living HeLa cells transfected with siRNA specific for indicated AP2 genes. (*lower*) Normalised NanoBiT assay results at 20 minutes post- addition of Nano-Glo live cell assay substrate (*n* = 5). Western blot analysis of AP2α1 and AP2μ2 protein in HeLa cells transfected with indicated siRNA. Data presented as mean ± S.E.M., *n* = 3-7, one- way ANOVA followed by Dunnett’s post hoc test (ns, not significant; \**P* < 0.05; \*\**P* < 0.01; \*\*\**P* < 0.001) or unpaired two-tailed t-test (^###^*P* < 0.001).

Despite a lack of evidence on how NIS endocytosis is controlled, we have previously identified the proto-oncogene PBF to be a NIS-interacting protein, capable of inducing NIS endocytosis via a canonical YXXΦ tyrosine-based endocytosis motif (174YARF177) (15). We next performed NanoBiT assays to determine whether the AP2 adaptor complex modulated the dynamic protein: protein interaction between NIS and PBF in living cells (**Fig. 1F**). First, we confirmed that exogenous PBF binds to NIS via NanoBiT assays (**Fig. 1G**), supporting our previous co-immunoprecipitation data (14). Critically, transient knockdown of the AP2 subunits α1 and μ2 markedly increased NIS binding to PBF in HeLa cells (*P* < 0.001; **Fig. 1G**), whereas transient inhibition of AP2σ2 expression had a marginal impact (Supp Fig. S2E). In support, AP2 knockdown with 4 distinct siRNAs enhanced NIS: PBF interaction in HEK293 cells (Supp Fig. S2F).

Together these data suggest that AP2-mediated functionality is key to controlling NIS expression and activity. Additionally, our findings imply that specific AP2 subunits alter the interaction of NIS with PBF, a key regulator of NIS activity, in living cells.

### Dissecting the mechanism of NIS interaction with the AP2 complex

NIS lacks a classical endocytosis motif (7,14). We therefore examined the intracellular C-terminus of NIS for motifs which might interact with the AP2 endocytic machinery and identified a potential dileucine (562**<u>LL</u>**563) and diacidic motif (578**<u>EE</u>**VA**<u>IL</u>**583), which bore clear structural similarity to the well characterised endocytosis motif of VMAT2 (**<u>EE</u>**KMA**<u>IL)</u>** (31). Species comparison revealed a high degree of conservation at the amino acid level of L and E residues (**Fig. 2A**). Abrogation of both motifs separately (L562A/L563A – *dileucine mutant,* and E578A/E579A – *diacidic mutant*) impacted NIS function. Whereas the dileucine mutant showed reduced RAI uptake ability, the diacidic mutant gained function (**Fig. 2B**). Biochemically, the dileucine mutant was partially glycosylated (**Fig. 2C**), and expressed in a general intracellular manner, in contrast to WT NIS, which is fully glycosylated and showed clear localisation at the PM, as well as intracellularly (**Fig. 2D**; Supp Fig. S2G and S2H). Compared to WT NIS, the diacidic mutant appeared to be completely glycosylated, with hypo- and mature glycosylated forms (**Fig. 2C**) but showed enhanced accumulation at the PM (**Fig. 2D**; Supp Fig. S2G and S2H).

**Figure 2.**
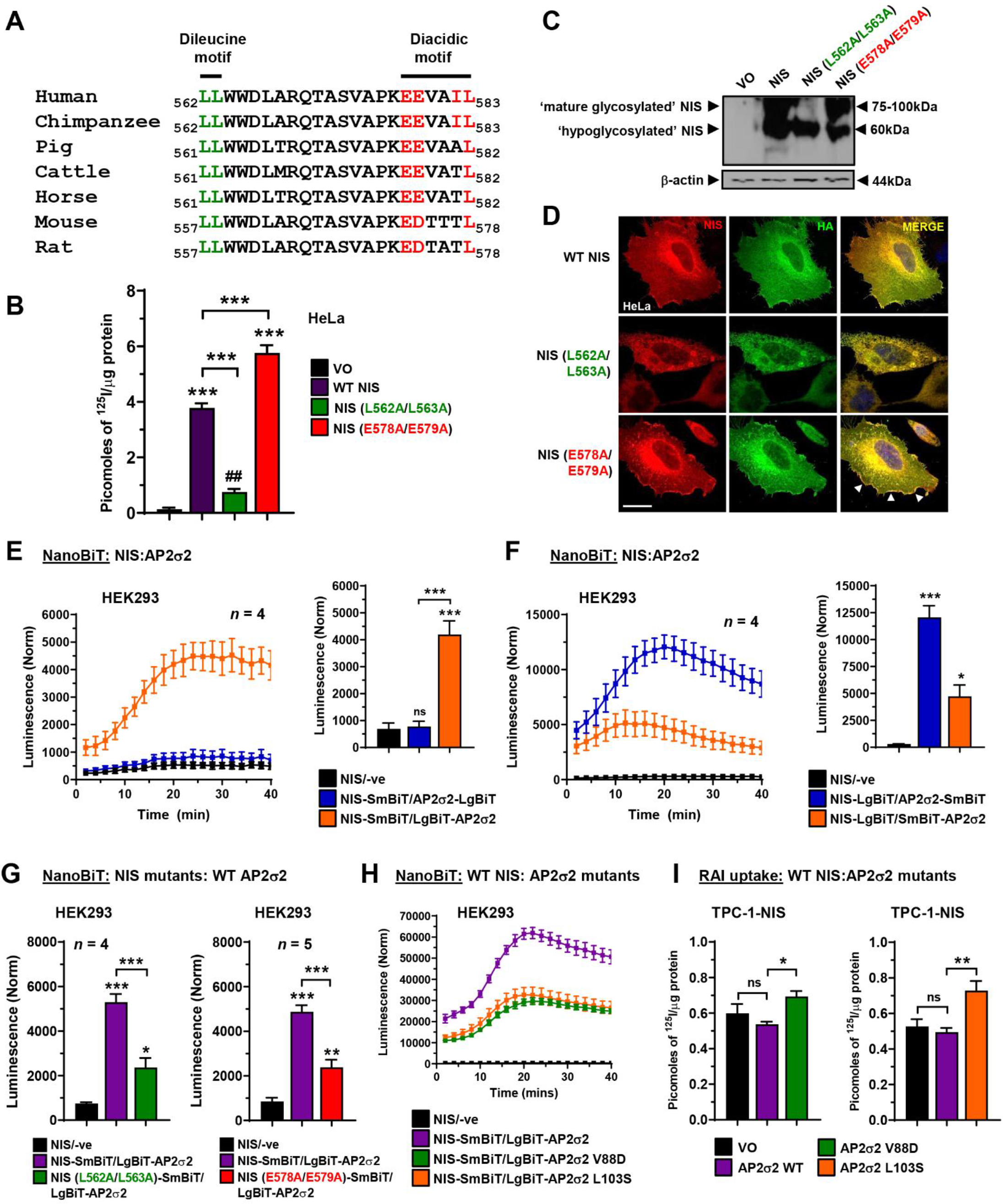
C-terminal motifs in NIS influence binding to AP2σ2 and are critical for function. **A,** Alignment of NIS C-terminus amino acid sequence (562–583) across multiple species. Potential dileucine (green) and diacidic (red) motifs are highlighted. **B,** RAI uptake in HeLa cells transfected with wild-type (WT) NIS, NIS mutant L562/L563A or NIS mutant E578A/E579A. **C,** Western blot analysis of different glycosylated isoforms of NIS protein in HeLa cells transfected with WT NIS, NIS mutant L562A/L563A and NIS mutant E578A/E579A. **D,** Same as **B** but confocal imaging in HeLa cells transfected with WT NIS and NIS mutants. Confocal images represent NIS expression (red), HA expression (green) and a merged image (yellow). Arrows (white) highlight PM regions with greater NIS localisation. Scale bar – 20 μm. See also Supp Fig. S1F and S1G. **E,** Live cell kinetic measurement using the NanoBiT assay to evaluate protein-protein interactions between NIS and AP2σ2 tagged with LgBiT in HEK293 cells. (*right*) NanoBiT assay results at 20 minutes post-addition of Nano-Glo substrate. See also Supp Fig. S2A. **F,** Same as **E** but AP2σ2 tagged with SmBiT. See also Supp Fig. S2B. **G,** Same as **E** but with NIS mutants L562A/L563A (*left*) and E578A/E579A (*right*). See also Supp Fig. S2C. **H,** Same as **E** but with AP2σ2 mutants V88D and L103S. See also Supp Fig. S2D and S2E. **I,** RAI uptake in TPC-1-NIS cells transfected with WT AP2σ2, AP2σ2 mutant V88D and AP2σ2 mutant L103S. See also Supp Fig. S2F. Data presented as mean ± S.E.M., *n* = 4-5, one-way ANOVA followed by Tukey’s post hoc test (ns, not significant; \**P* < 0.05; \*\**P* < 0.01; \*\*\**P* < 0.001) or unpaired two-tailed t-test (^##^*P* < 0.01).

Given that the σ2 subunit of AP2 is known to bind diacidic/dileucine motifs of proteins prior to initiating endocytosis (32), we next assessed the impact of the dileucine and diacidic mutants on binding to the AP2σ2 subunit. After subcloning the σ2 cDNA, we detected binding to NIS in NanoBiT assays; WT NIS SmBiT bound to N-terminal LgBiT σ2 (**Fig. 2E**; Supp Fig. S3A), as well as NIS LgBiT binding to σ2 tagged with SmBiT on both the N and C-termini (**Fig. 2F**; Supp Fig. S3B). Of particular significance, the diacidic and dileucine mutants both bound σ2 with reduced stringency compared to WT NIS in live cells (**Fig. 2G**; Supp Fig. S3C). We next investigated whether any regions of σ2 might be important for interaction with NIS. Based on previous observations identifying key σ2 residues for interacting with diacidic/dileucine motifs (32), we found that V88D and L103S σ2 mutants gave reduced binding to NIS via NanoBiT assays in HEK293 and HeLa cells (**Fig. 2H**; Supp Fig. S3D and S3E). In support of this, each AP2σ2 mutation led to greater RAI uptake in thyroidal TPC-1-NIS and 8505C-NIS cells compared to WT AP2σ2 (**Fig. 2I**; Supp Fig. S3F). Thus, we detected NIS binding to AP2σ2 in live cells, albeit with different efficiencies depending on N- or C-terminal tagging. Mutation of the diacidic and dileucine motifs reduced NIS binding to AP2σ2, presumably due at least in part to reduced PM localisation for the dileucine mutation. AP2σ2 residues V88 and L103 appear important to NIS interaction, and we hypothesise that over-expression of σ2 subunits with reduced capacity to bind NIS competitively inhibits endogenous WT σ2 function within the AP2 heterotetramer, resulting in NIS accumulation at the PM and hence increased RAI uptake.

### AP2α genes are associated with recurrence in RAI-treated patients

Having identified AP2 genes as key NIS regulators, we next appraised TCGA and GEO datasets to investigate their clinical relevance and prognostic utility. Of significance, *AP2A1*, *AP2B1* and *AP2M1* were highly expressed in papillary thyroid cancer (PTC) with the more aggressive BRAF-like gene signature compared to RAS-like PTC (**Fig. 3A**), as well as in the independent PTC dataset GSE60542 (**Fig. 3B**). AP2A1 expression was also greater in recurrent PTC (THCA; *n* = 486), as well as in RAI-treated PTC (*n* = 256) (**Fig. 3C**). ROC determination of optimal cut-offs in the RAI-treated BRAF-like PTC cohort (**Fig. 3D**) revealed a significant reduction in DFS associated with higher expression of AP2A1 and AP2A2 in PTC (**Fig. 3E**; Supp Fig. S4A), as well as an increased risk of recurrence (**Table 1A**). In particular, the risk of recurrence with high AP2A2 was greatest for RAI-treated patients associated with *BRAF* mutations (**Table 1A**). Multivariate analysis further showed that AP2A2 was an independent predictive factor for recurrence in BRAF mutated RAI-treated PTC after controlling for multiple clinical variables and all five AP2 genes (HR 6.310, 95% CI 1.695-24.962; **Table 2A**). Importantly, AP2 gene expression lacked any strong association with cancer staging in BRAF mutated RAI-treated PTC (Supp Fig. S4B), suggesting that any predictive value was likely related to a poorer response to RAI therapy rather than tumour aggressive features. In support, there was no association between cancer staging and AP2 genes in the independent PTC cohort GSE60542 (Supp Fig. S4C and S4D).

**Figure 3.**
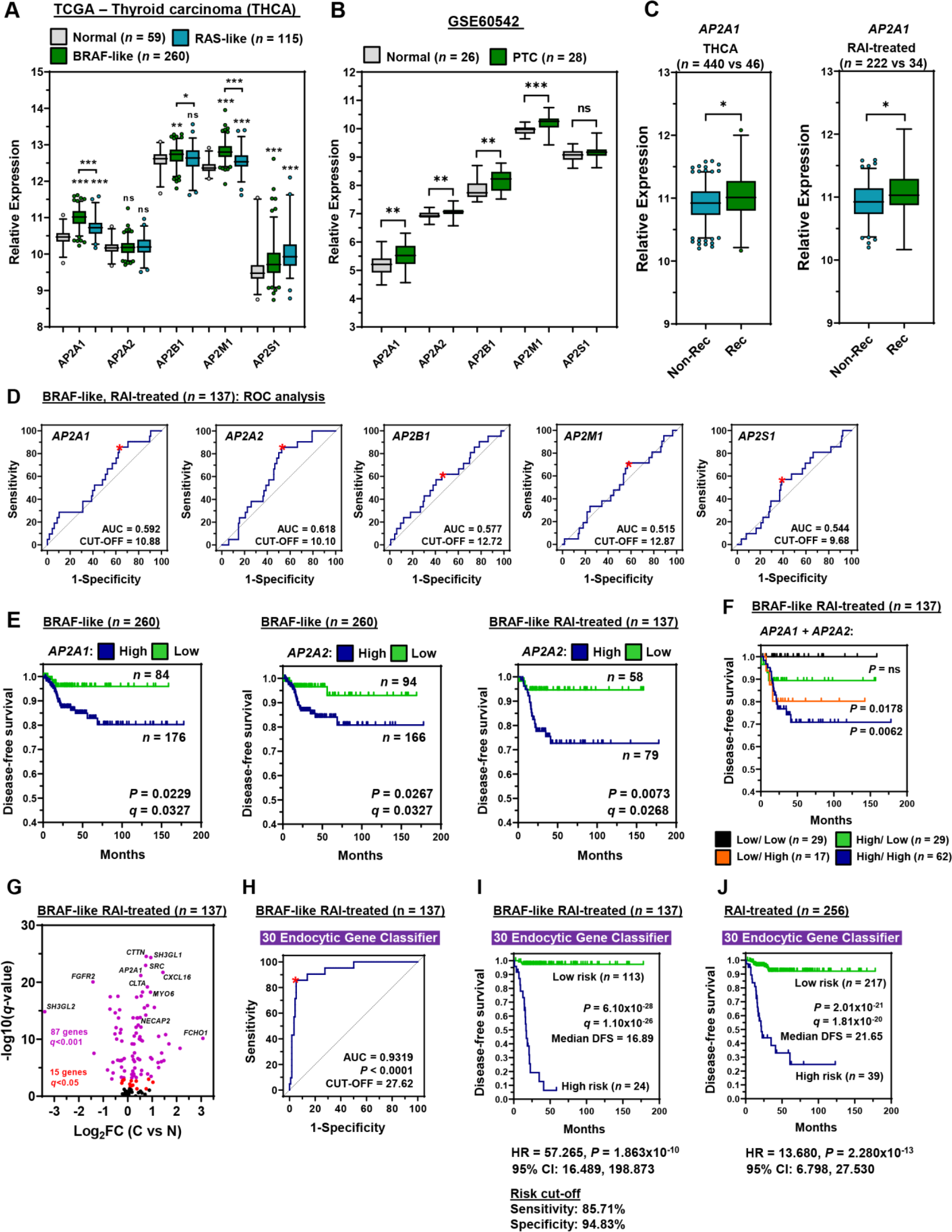
AP2 gene-related risk score classifier is predictive of thyroid cancer recurrence. **A** and **B,** Box and whisker plots showing expression (log2) of AP2 genes in the (**A**) THCA (BRAF-like and RAS-like PTC versus normal) and (**B**) GSE60542 (PTC versus normal) datasets. **C,** Box and whisker plots showing AP2A1 expression in the THCA (*left*) and RAI-treated (*right*) cohorts [recurrent (REC) versus non- recurrent (NON-REC)]. **D,** Representative ROC curves of AP2 genes in the BRAF-like, RAI-treated THCA cohort (*n* = 137). **E,** Representative Kaplan-Meier analysis of DFS for the BRAF-like and BRAF-like, RAI treated THCA cohorts stratified on high versus low tumoral expression of indicated AP2 genes; log-rank test. Number (*n*) of patients per expression sub-group (high/low), *P*-values and *q*- values are shown. **F,** Same as **E** but patients stratified on high versus low tumoral expression for both AP2A1 and AP2A2 in the BRAF-like, RAI treated THCA cohort. **G,** Volcano plot comparing log2FC with *q*-value (-log base 10) for the BRAF-like, RAI-treated THCA cohort [C versus N; *n* = 137] and 137 endocytosis-related genes. See also Supp Fig. S4A. **H-I,** ROC analysis (**H**) and Kaplan-Meier curve (**I**) of the 30 endocytic gene risk score classifier in the BRAF-like, RAI-treated THCA cohort. **J,** Same as **I** but with the RAI-treated THCA cohort (*n* = 256). *P*-values were determined by Mann-Whitney test and adjusted using the Benjamini-Hochberg FDR correction procedure (ns, not significant; \**P* < 0.05; \*\**P* < 0.01; \*\*\**P* < 0.001).

**Table 1.**
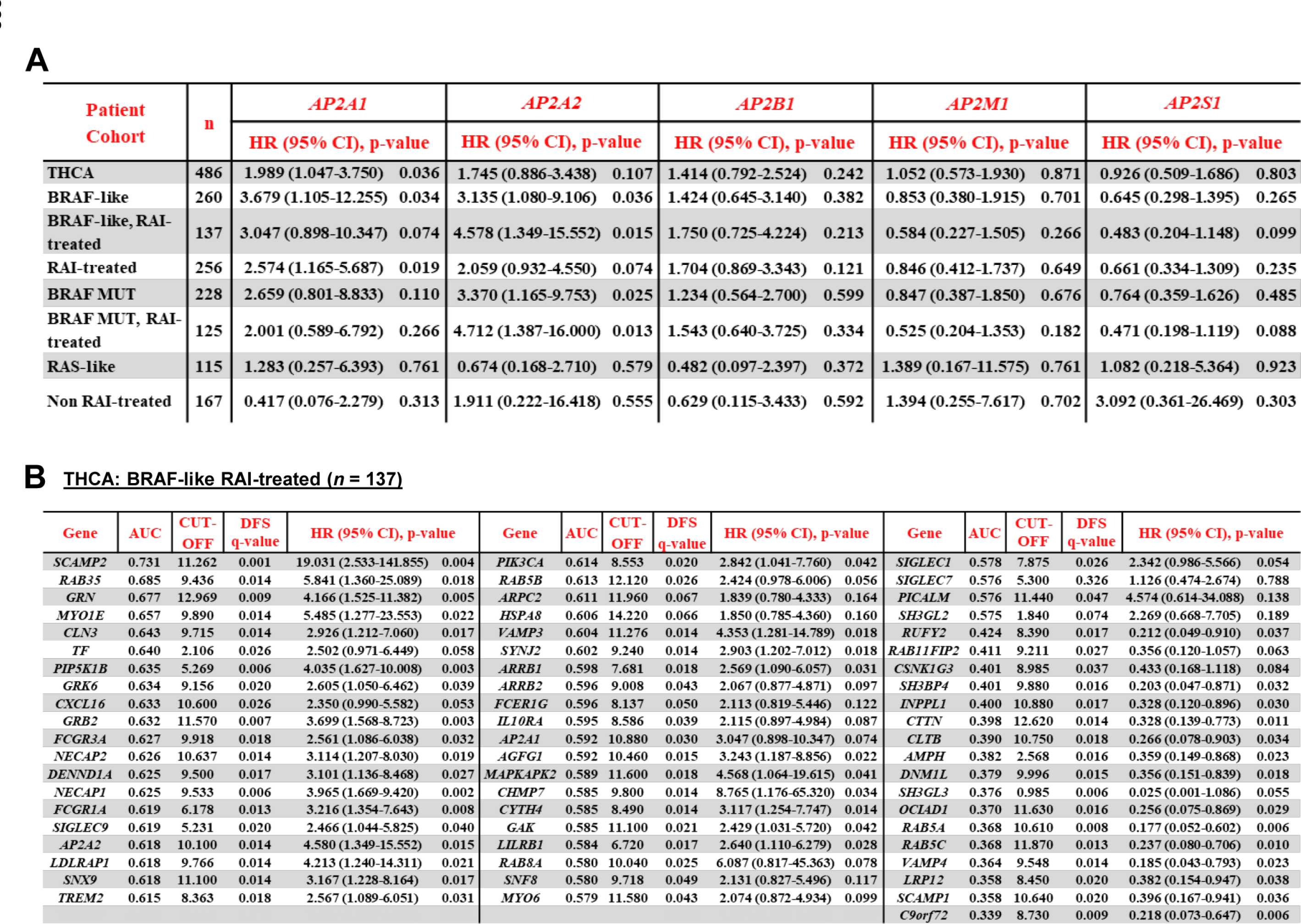
ROC and Cox regression analysis of endocytosis genes. **A,** Univariate Cox regression analysis in different THCA patient cohorts comparing AP2 genes. *n*, number; HR, hazard ratio; CI, confidence interval. **B,** ROC and univariate Cox regression analysis of 61 endocytosis-related genes in BRAF-like, RAI treated THCA (*n* = 137). AUC, area under curve; CUT-OFF, cut-off expression values used for patient cohort stratification; DFS, disease-free survival; HR, hazard ratio; CI, confidence interval.

**Table 2.**
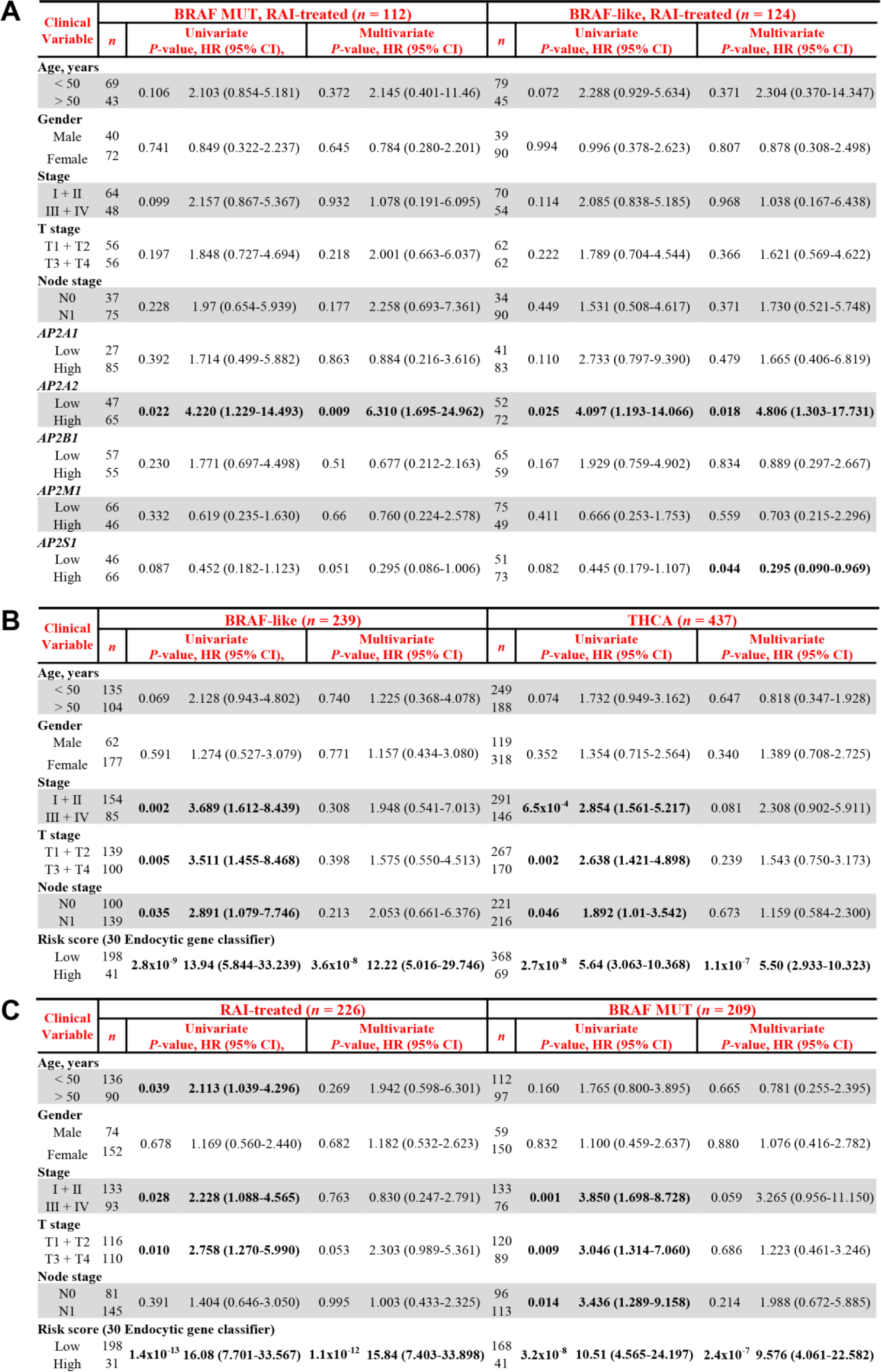
Multivariate analysis of THCA patient cohorts. **A,** *n*, number; HR. hazard ratio; CI, confidence interval. *P*-values in bold were less than 0.05 and considered statistically significant. Some patients in the BRAF MUT, RAI treated (*n* = 13) and BRAF-like, RAI-treated (*n* = 13) cohorts were not included in univariate and multivariate analysis due to missing clinical variables. **B** and **C**, Same as **A** except the 30 endocytic gene risk score classifier was used instead of the individual AP2 genes in larger THCA cohorts as indicated. Some patients in the BRAF-like (*n* = 21), THCA (*n* = 49), RAI-treated (*n* = 30) and BRAF MUT (*n* = 19) cohorts were not included in univariate and multivariate analysis due to missing clinical variables.

| Clinical Variable | BRAF MUT, RAI-treated (n = 112) |  |  |  |  | BRAF-like, RAI-treated (n = 124) |  |  |  |  |
| --- | --- | --- | --- | --- | --- | --- | --- | --- | --- | --- |
|  | n | Univariate<br>P-value, HR (95% CI), |  | Multivariate<br>P-value, HR (95% CI) |  | n | Univariate<br>P-value, HR (95% CI) |  | Multivariate<br>P-value, HR (95% CI) |  |
| Age, years |  |  |  |  |  |  |  |  |  |  |
| < 50 | 69 |  |  |  |  | 79 |  |  |  |  |
| > 50 | 43 | 0.106 | 2.103 (0.854-5.181) | 0.372 | 2.145 (0.401-11.46) | 45 | 0.072 | 2.288 (0.929-5.634) | 0.371 | 2.304 (0.370-14.347) |
| Gender |  |  |  |  |  |  |  |  |  |  |
| Male | 40 |  |  |  |  | 39 |  |  |  |  |
| Female | 72 | 0.741 | 0.849 (0.322-2.237) | 0.645 | 0.784 (0.280-2.201) | 90 | 0.994 | 0.996 (0.378-2.623) | 0.807 | 0.878 (0.308-2.498) |
| Stage |  |  |  |  |  |  |  |  |  |  |
| I + II | 64 |  |  |  |  | 70 |  |  |  |  |
| III + IV | 48 | 0.099 | 2.157 (0.867-5.367) | 0.932 | 1.078 (0.191-6.095) | 54 | 0.114 | 2.085 (0.838-5.185) | 0.968 | 1.038 (0.167-6.438) |
| T stage |  |  |  |  |  |  |  |  |  |  |
| T1 + T2 | 56 |  |  |  |  | 62 |  |  |  |  |
| T3 + T4 | 56 | 0.197 | 1.848 (0.727-4.694) | 0.218 | 2.001 (0.663-6.037) | 62 | 0.222 | 1.789 (0.704-4.544) | 0.366 | 1.621 (0.569-4.622) |
| Node stage |  |  |  |  |  |  |  |  |  |  |
| N0 | 37 |  |  |  |  | 34 |  |  |  |  |
| N1 | 75 | 0.228 | 1.97 (0.654-5.939) | 0.177 | 2.258 (0.693-7.361) | 90 | 0.449 | 1.531 (0.508-4.617) | 0.371 | 1.730 (0.521-5.748) |
| AP2A1 |  |  |  |  |  |  |  |  |  |  |
| Low | 27 |  |  |  |  | 41 |  |  |  |  |
| High | 85 | 0.392 | 1.714 (0.499-5.882) | 0.863 | 0.884 (0.216-3.616) | 83 | 0.110 | 2.733 (0.797-9.390) | 0.479 | 1.665 (0.406-6.819) |
| AP2A2 |  |  |  |  |  |  |  |  |  |  |
| Low | 47 |  |  |  |  | 52 |  |  |  |  |
| High | 65 | **0.022** | **4.220 (1.229-14.493)** | **0.009** | **6.310 (1.695-24.962)** | 72 | **0.025** | **4.097 (1.193-14.066)** | **0.018** | **4.806 (1.303-17.731)** |
| AP2B1 |  |  |  |  |  |  |  |  |  |  |
| Low | 57 |  |  |  |  | 65 |  |  |  |  |
| High | 55 | 0.230 | 1.771 (0.697-4.498) | 0.51 | 0.677 (0.212-2.163) | 59 | 0.167 | 1.929 (0.759-4.902) | 0.834 | 0.889 (0.297-2.667) |
| AP2M1 |  |  |  |  |  |  |  |  |  |  |
| Low | 66 |  |  |  |  | 75 |  |  |  |  |
| High | 46 | 0.332 | 0.619 (0.235-1.630) | 0.66 | 0.760 (0.224-2.578) | 49 | 0.411 | 0.666 (0.253-1.753) | 0.559 | 0.703 (0.215-2.296) |
| AP2S1 |  |  |  |  |  |  |  |  |  |  |
| Low | 46 |  |  |  |  | 51 |  |  |  |  |
| High | 66 | 0.087 | 0.452 (0.182-1.123) | 0.051 | 0.295 (0.086-1.006) | 73 | 0.082 | 0.445 (0.179-1.107) | **0.044** | **0.295 (0.090-0.969)** |

B
| Clinical Variable | BRAF-like (*n* = 239) |  |  |  |  | THCA (*n* = 437) |  |  |  |  |
| --- | --- | --- | --- | --- | --- | --- | --- | --- | --- | --- |
|  | *n* | Univariate<br>*P*-value, HR (95% CI), |  | Multivariate<br>*P*-value, HR (95% CI) |  | *n* | Univariate<br>*P*-value, HR (95% CI) |  | Multivariate<br>*P*-value, HR (95% CI) |  |
| Age, years |  |  |  |  |  |  |  |  |  |  |
| < 50 | 135 |  |  |  |  | 249 |  |  |  |  |
| > 50 | 104 | 0.069 | 2.128 (0.943-4.802) | 0.740 | 1.225 (0.368-4.078) | 188 | 0.074 | 1.732 (0.949-3.162) | 0.647 | 0.818 (0.347-1.928) |
| Gender |  |  |  |  |  |  |  |  |  |  |
| Male | 62 |  |  |  |  | 119 |  |  |  |  |
| Female | 177 | 0.591 | 1.274 (0.527-3.079) | 0.771 | 1.157 (0.434-3.080) | 318 | 0.352 | 1.354 (0.715-2.564) | 0.340 | 1.389 (0.708-2.725) |
| Stage |  |  |  |  |  |  |  |  |  |  |
| I + II | 154 | 0.002 | 3.689 (1.612-8.439) | 0.308 | 1.948 (0.541-7.013) | 291 | 6.5x10<sup>-4</sup> | 2.854 (1.561-5.217) | 0.081 | 2.308 (0.902-5.911) |
| III + IV | 85 |  |  |  |  | 146 |  |  |  |  |
| T stage |  |  |  |  |  |  |  |  |  |  |
| T1 + T2 | 139 | 0.005 | 3.511 (1.455-8.468) | 0.398 | 1.575 (0.550-4.513) | 267 | 0.002 | 2.638 (1.421-4.898) | 0.239 | 1.543 (0.750-3.173) |
| T3 + T4 | 100 |  |  |  |  | 170 |  |  |  |  |
| Node stage |  |  |  |  |  |  |  |  |  |  |
| N0 | 100 | 0.035 | 2.891 (1.079-7.746) | 0.213 | 2.053 (0.661-6.376) | 221 | 0.046 | 1.892 (1.01-3.542) | 0.673 | 1.159 (0.584-2.300) |
| N1 | 139 |  |  |  |  | 216 |  |  |  |  |
| Risk score (30 Endocytic gene classifier) |  |  |  |  |  |  |  |  |  |  |
| Low | 198 | 2.8x10<sup>-9</sup> | 13.94 (5.844-33.239) | 3.6x10<sup>-8</sup> | 12.22 (5.016-29.746) | 368 | 2.7x10<sup>-8</sup> | 5.64 (3.063-10.368) | 1.1x10<sup>-7</sup> | 5.50 (2.933-10.323) |
| High | 41 |  |  |  |  | 69 |  |  |  |  |

**Table 2.**
| Clinical Variable | RAI-treated (n = 226) |  |  |  |  | BRAF MUT (n = 209) |  |  |  |  |
| --- | --- | --- | --- | --- | --- | --- | --- | --- | --- | --- |
|  | n | Univariate<br>P-value, HR (95% CI), |  | Multivariate<br>P-value, HR (95% CI) |  | n | Univariate<br>P-value, HR (95% CI) |  | Multivariate<br>P-value, HR (95% CI) |  |
| Age, years |  |  |  |  |  |  |  |  |  |  |
| < 50 | 136 |  |  |  |  | 112 |  |  |  |  |
| > 50 | 90 | 0.039 | 2.113 (1.039-4.296) | 0.269 | 1.942 (0.598-6.301) | 97 | 0.160 | 1.765 (0.800-3.895) | 0.665 | 0.781 (0.255-2.395) |
| Gender |  |  |  |  |  |  |  |  |  |  |
| Male | 74 |  |  |  |  | 59 |  |  |  |  |
| Female | 152 | 0.678 | 1.169 (0.560-2.440) | 0.682 | 1.182 (0.532-2.623) | 150 | 0.832 | 1.100 (0.459-2.637) | 0.880 | 1.076 (0.416-2.782) |
| Stage |  |  |  |  |  |  |  |  |  |  |
| I + II | 133 |  |  |  |  | 133 |  |  |  |  |
| III + IV | 93 | 0.028 | 2.228 (1.088-4.565) | 0.763 | 0.830 (0.247-2.791) | 76 | 0.001 | 3.850 (1.698-8.728) | 0.059 | 3.265 (0.956-11.150) |
| T stage |  |  |  |  |  |  |  |  |  |  |
| T1 + T2 | 116 |  |  |  |  | 120 |  |  |  |  |
| T3 + T4 | 110 | 0.010 | 2.758 (1.270-5.990) | 0.053 | 2.303 (0.989-5.361) | 89 | 0.009 | 3.046 (1.314-7.060) | 0.686 | 1.223 (0.461-3.246) |
| Node stage |  |  |  |  |  |  |  |  |  |  |
| N0 | 81 |  |  |  |  | 96 |  |  |  |  |
| N1 | 145 | 0.391 | 1.404 (0.646-3.050) | 0.995 | 1.003 (0.433-2.325) | 113 | 0.014 | 3.436 (1.289-9.158) | 0.214 | 1.988 (0.672-5.885) |
| Risk score (30 Endocytic gene classifier) |  |  |  |  |  |  |  |  |  |  |
| Low | 198 |  |  |  |  | 168 |  |  |  |  |
| High | 31 | 1.4x10<sup>-13</sup> | 16.08 (7.701-33.567) | 1.1x10<sup>-12</sup> | 15.84 (7.403-33.898) | 41 | 3.2x10<sup>-8</sup> | 10.51 (4.565-24.197) | 2.4x10<sup>-7</sup> | 9.576 (4.061-22.582) |

The AP2α subunit has 2 major isoforms encoded by 2 separate genes, *AP2A1* and *AP2A2*; we investigated the differential expression of both *AP2A1* and *AP2A2* to fully define the prognostic utility of AP2α. Importantly, clinical data showed poorer DFS for patients with high tumoural AP2A1/AP2A2 than for other patient groups (**Fig. 3F**; Supp Fig. S4E). There was also greater recurrence for patients with high AP2A1/AP2A2 than those stratified on AP2A1 or AP2A2 combined with other AP2 genes (Supp Fig. S4F). These findings indicate that the status of both AP2α genes should be regarded as an important clinical indicator for recurrence, especially in RAI-treated patients.

### Endocytic genes are independent predictive indicators of recurrence

We next challenged the BRAF-like, RAI-treated PTC transcriptome against *AP2A2* expression to better understand prominent biological pathways associated with recurrence (Supp Fig. S5A and S5B). In support of our findings, functional analyses (DAVID, ToppGene) revealed endocytosis and protein transport as key dysregulated pathways (Supp Fig. S5C and S5D), as well as identifying 102 endocytosis-related genes with differential expression (C versus N; **Fig. 3G**). Hierarchical cluster analysis of the 61 most clinically relevant endocytic genes (Supp Fig 5E; **Table 1B**) revealed 4 major patient clusters (Supp Fig. 6A). Of particular significance, patients associated with recurrence (clusters 2 and 4; Supp Fig. S6B) had greater endocytic gene dysregulation (Supp Fig. S6C), and higher expression of AP2A1 and AP2A2 (Supp Fig. S6D-F).

We next evaluated a panel of multigene risk score classifiers for predicting recurrence based on endocytic genes associated with highest recurrence (subcluster 4a versus clusters 1 and 3; Supp Fig. S6G, S7A; Supp Table S2). Importantly, a higher AUC of 0.9319 (**Fig. 3H**) indicated a greater prediction effect for the 30 endocytic gene-based risk score compared to individual genes (AUC 0.575-0.731; **Table 1B**). In agreement, there was a significant association with poorer DFS in BRAF-like, RAI-treated PTC (median DFS = 16.89 months; **Fig. 3I**). Patients at higher risk also had a significantly worse prognosis (HR = 57.265, 95% CI 16.489-198.873; **Fig. 3I**), which was validated in larger THCA cohorts (**Fig. 3J**; Supp Fig. S7B-S7D). By contrast, there was no predictive effect in non-RAI treated or RAS-like THCA (Supp Fig. S7C and S7D). Critically, multivariate analysis further showed that the 30 gene risk score was an independent predictive factor for larger THCA cohorts (**Table 2B and C**).

### Manipulating endocytosis to enhance NIS function

Given our evidence of extensive dysregulation of endocytic genes and association with recurrence, we next appraised whether endocytosis can be exploited as a druggable strategy to enhance RAI uptake. Endocytosis inhibitors already exist (33) but there is no known chemical means of directly modulating NIS endocytosis. Utilizing NIS in high-throughput drug screening we recently identified that the anti-malarial drug chloroquine (CQ) significantly induced RAI uptake, peaking at 8 hr post-treatment (23). Here, we investigated whether CQ might act upon NIS endocytosis, as a rapid mechanism of functional modulation, and might therefore represent the first pharmaceutical agent for altering NIS endocytosis. A significant finding was that siRNA ablation of the AP2α1 subunit blocked CQ’s induction of RAI uptake in human primary thyrocytes (**Fig. 4A**). In addition, CQ was unable to induce significant RAI uptake in TPC-1-NIS and 8505C-NIS cells when AP2α1 was abrogated (**Fig. 4B and 4C**).

**Figure 4.**
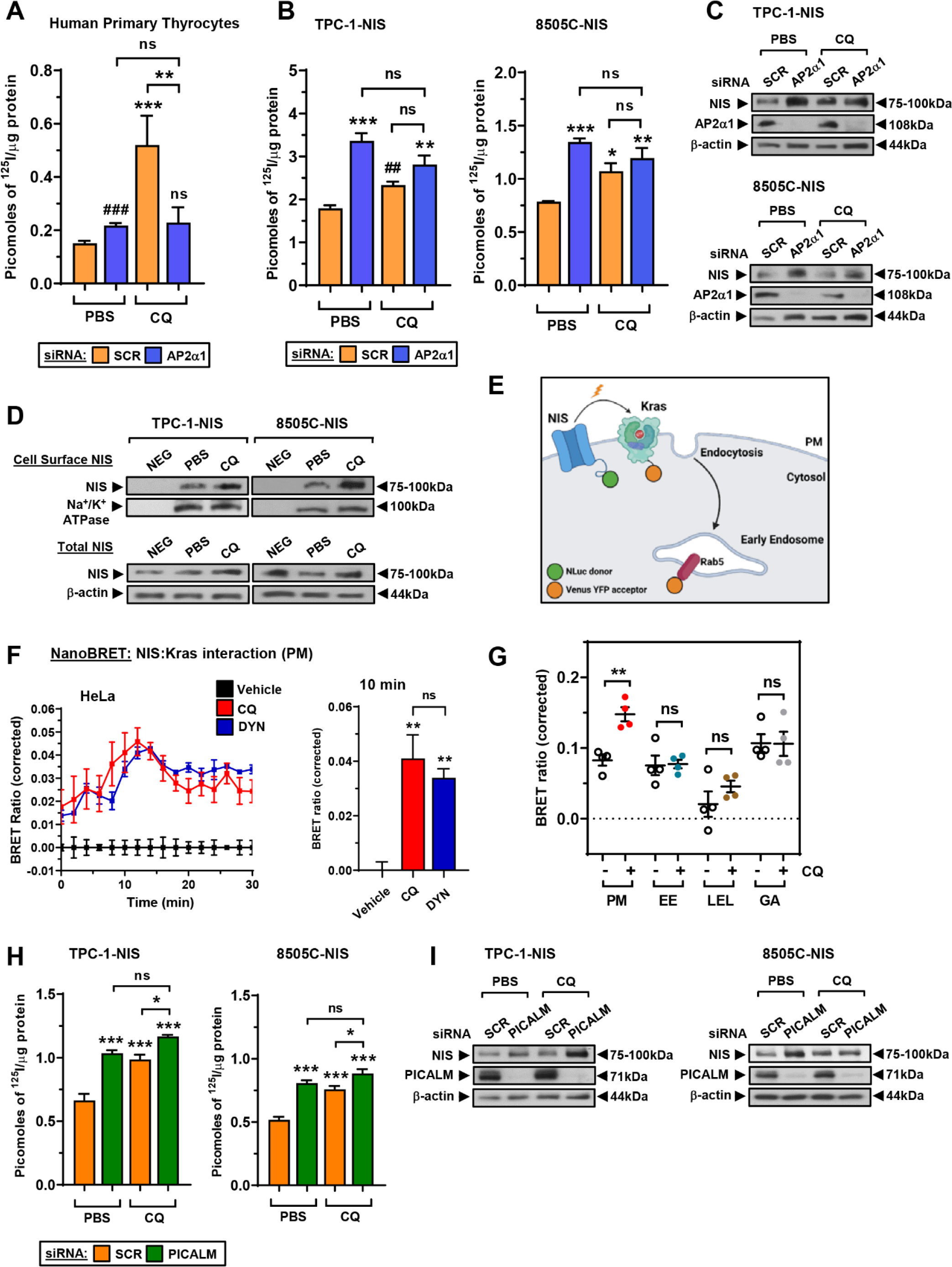
CQ inhibits endocytosis to increase NIS protein at the plasma membrane. **A,** RAI uptake in human primary thyrocytes following AP2α1-siRNA depletion and chloroquine (CQ) treatment. Scr – scrambled control siRNA. **B** and **C,** RAI uptake (**B**) and relative NIS and AP2α1 protein levels (**C**) in TPC-1-NIS and 8505C-NIS cells following AP2α1-siRNA depletion and CQ treatment. Scr – scrambled control siRNA. **D,** Western blot analysis of NIS protein at the PM relative to Na^+^/K^+^ ATPase following CSBA in TPC-1-NIS and 8505C-NIS cells after CQ treatment. (*lower*) Total NIS protein levels in thyroid cells treated with CQ. **E,** Schematic depicting NanoBRET assay to monitor close proximity of NIS with highly abundant PM proteins (e.g. Kras). Created with BioRender.com. **F,** Live cell kinetic measurement using the NanoBRET signal to evaluate the close proximity between NIS and Kras in HeLa cells treated with CQ or DYN. (*right*) NanoBRET assay results at 10 minutes post- addition of Nano-Glo substrate. **G,** Profiling PM and subcellular changes of NIS using the NanoBRET assay in CQ-treated HeLa cells. HeLa cells were transiently transfected with NIS tagged with NLuc, and the PM marker Kras or one of the subcellular markers Rab5 (EE, early endosome), Rab7 (LEL, late endosome/lysosome) or Rab6 (GA, golgi apparatus) tagged with Venus. **H** and **I,** RAI uptake (**H**) and relative NIS and PICALM protein levels (**I**) in TPC-1-NIS and 8505C-NIS cells following PICALM- siRNA depletion and CQ treatment. Scr – scrambled control siRNA.Data presented as mean ± S.E.M., *n* = 3-4, one-way ANOVA followed by Tukey’s post hoc test (ns, not significant; \**P* < 0.05; \*\**P* < 0.01; \*\*\**P* < 0.001) or unpaired two-tailed t-test (^#^*P* < 0.05; ^##^*P* < 0.01).

To better understand how CQ influences NIS function, we characterised NIS expression at the PM via cell surface biotinylation assays (CSBA), which demonstrated elevated levels of cell-surface NIS in CQ-treated thyroid cancer cells (**Fig. 4D**). This finding was confirmed in live cells via NanoBRET assays (**Fig. 4E**), in which a BRET signal is generated when NIS is in close proximity with the abundant PM protein Kras. Critically, CQ gave a strong BRET signal similar to Dynasore treatment of cells (**Fig. 4F**), which occurred predominately at the PM rather than at other intracellular locations (**Fig. 4G**, Supp Fig. S8A and S8B). As a control, Dynasore had a potent impact on RAI uptake (Supp Fig. S8C). To challenge our findings, we next investigated whether ablation of the endocytic factor PICALM, identified as a putative NIS interactor (**Fig. 1A**) and known to recruit AP2/clathrin to the PM (34), would also affect the ability of CQ to enhance RAI uptake. The abrogation of PICALM significantly enhanced NIS expression and function in a similar manner to AP2α ablation, as well as blunting the induction of RAI uptake by CQ in thyroid cancer cells (**Fig. 4H and 4I**).

Overall we thus hypothesise that CQ’s induction of radioiodide uptake reflects its interference with the PICALM/AP2/clathrin machinery which controls NIS endocytosis.

### CQ and SAHA enhance thyroidal ^99m^Tc pertechnetate uptake in mice

We recently demonstrated that combining drugs with distinct modes of action, such as CQ with the HDAC inhibitor SAHA, gave robust and additive increases in RAI uptake in thyroid cells in vitro (23). Here, the combination of AP2α ablation and SAHA administration gave a similar robust and significant increase in RAI uptake in thyroid cancer cells compared to each treatment alone (Supp Fig. S8D), indicating that inhibiting endocytosis enhances the impact of SAHA on NIS function. We next progressed our approaches to WT BALB/c mice to examine the translatable potential of CQ with SAHA to improve endogenous NIS function (**Fig. 5A**). Importantly, co-treatment with CQ and SAHA led to a significant increase in thyroidal uptake of the radiotracer technetium-99m pertechnetate (^99m^Tc) after 30 min (52.7%; *P* = 0.0003; **Fig. 5B**) versus controls. By comparison, neither CQ nor SAHA had any impact on ^99m^Tc uptake in mice thyroid glands when administered separately (*P* = NS; **Fig. 5B**). In addition, CQ+SAHA induced NIS mRNA in mouse thyroids (2.2-fold vs CON; *P* < 0.0001; **Fig. 5C**) at significantly higher levels than in CQ- (*P* < 0.001) or SAHA-treated BALB/c mice (*P* < 0.05).

**Figure 5.**
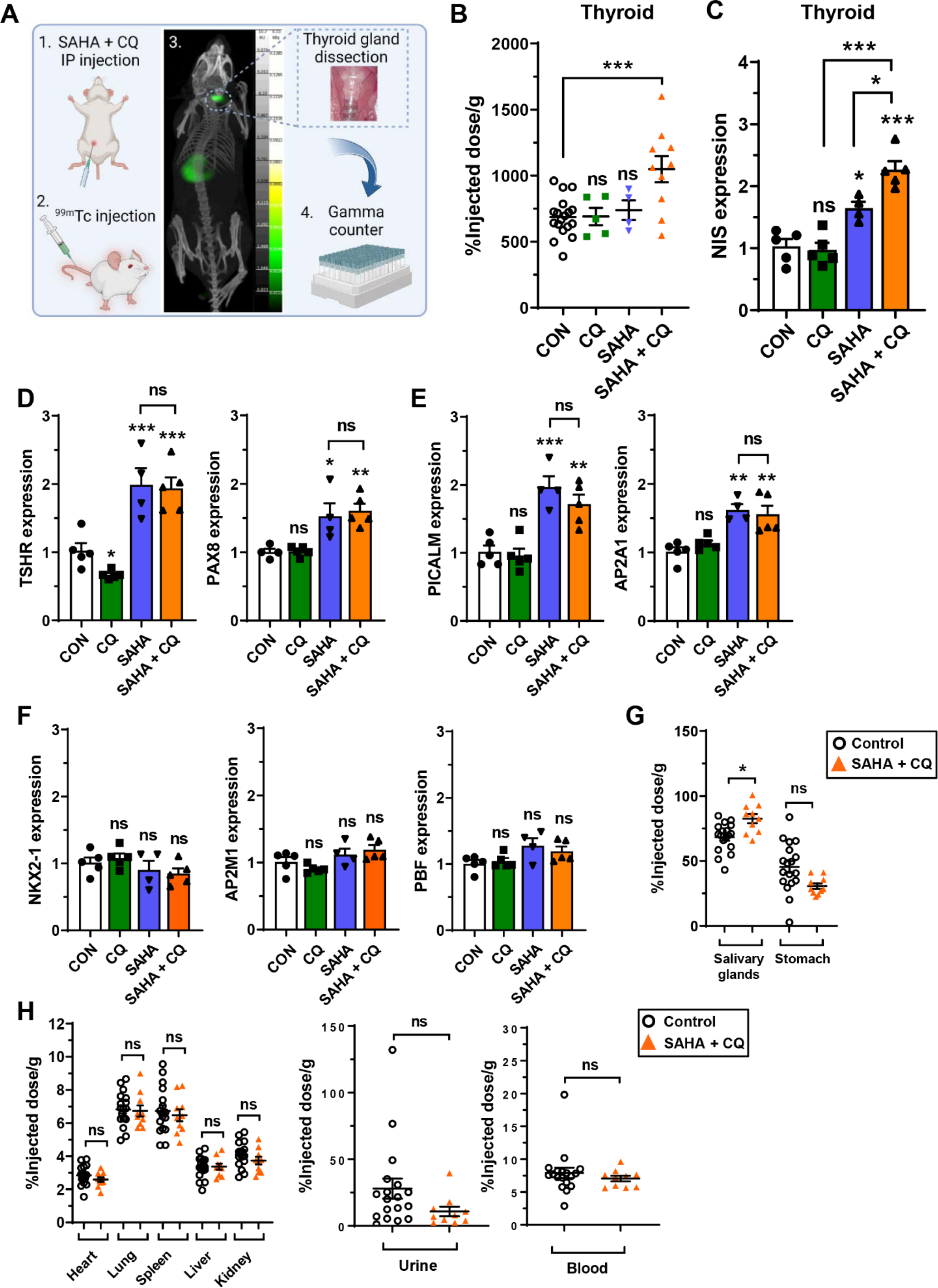
Targeting endocytosis to enhance the impact of SAHA on NIS function in vivo. **A,** Schematic of steps (1–4) used to examine the translatable potential of CQ and SAHA to enhance NIS function *in vivo*. Created with BioRender.com. **B** and **C**, Technetium-99m pertechnetate (^99m^Tc) uptake (**B**; *n* = 4 -18) and NIS mRNA levels (**C**) in thyroid glands dissected from WT BALB/c mice administered with CQ and SAHA either alone or in combination. **D-F,** Same as **C** but comparing relative TSHR, PAX8, PICALM, AP2A1, NKX2-1, AP2M1 and PBF mRNA levels in mouse thyroids. **G and H,** Distribution of ^99m^Tc uptake across the indicated tissues harvested from WT BALB/c mice treated as described in **B**. Data presented as mean ± S.E.M., *n* = 3-5, one-way ANOVA followed by Tukey’s post hoc test (ns, not significant; \**P* < 0.05; \*\**P* < 0.01; \*\*\**P* < 0.001).

NIS expression is regulated at multiple levels, including transcriptional and post-translational mechanisms (7). Here, we found that elevated thyroidal NIS mRNA in SAHA- and CQ+SAHA-treated mice was associated with higher expression of well-known NIS regulators, including TSHR and PAX8 (**Fig. 5D**). Interestingly, PICALM and AP2A1 were also induced in SAHA-treated mice (**Fig. 5E**). In contrast, there was no change in expression of Nkx2-1, PBF, AP2M1 or any controls (**Fig. 5F** and Supp Fig 8E). Subsequent biodistribution studies revealed a marginal increase in ^99m^Tc uptake in NIS-expressing salivary glands of CQ+SAHA-treated mice (20.9%; *P* < 0.05; **Fig. 5G**), but no differences in other tissues (**Fig. 5H**; Supp Fig. S8F), or any change in mice body weight (Supp Fig. S8G). In contrast to CQ, control experiments validated the impact of SAHA on NIS, TSHR, AP2A1 and PICALM mRNA in thyroid cancer cells (Supp Fig. S8H-J).

Together these findings demonstrate that the combination of CQ and SAHA enhances endogenous thyroidal NIS expression and function in vivo, highlighting the potential of pharmacologically inhibiting NIS endocytosis to increase therapeutic radionuclide uptake in patients with thyroid cancer.

## DISCUSSION

The sodium iodide symporter is the sole known conduit of iodide into human cells, and as such is exploited in the ablation of thyroid cancers and their metastases, as well as in various other clinical and pre-clinical settings (35,36). In enhancing the uptake of radionuclides via NIS it is critical to consider the mechanisms which underlie its movement into and away from the PM. Here, combining our previously published mass spectrometry data (11) with that of Faria et al (28), we identified AP2 complex genes to be NIS interactors in both independent studies, aligning with previous circumstantial evidence implicating the AP2 complex in clathrin-dependent endocytosis of NIS (14).

The heterotetrameric AP2 adaptor complex comprises two large subunits (α and β2), one medium subunit (μ2), and one small subunit (σ2) (37) and is the key effector of clathrin-dependent endocytosis, interacting directly with clathrin via its α and β subunits. Recent screening data revealed that abrogating the AP2μ2-subunit results in marked enrichment of the proto-oncogene PBF in the PM (9.6-fold enriched, 4^th^ highest of all identified proteins) (38), further implicating AP2μ2 in PBF function. The σ2 subunit of AP2 is known to bind diacidic/dileucine motifs of proteins, whereas the μ2 subunit of AP2 binds YXXΦ motifs (32,39). Summating our current and previous data, our current hypothesis is that the AP2 heterotetramer binds NIS in a stable conformation at the PM, mediated by direct interaction with σ2, ahead of a ‘final signal’ for endocytosis to progress, impacted by the μ2 subunit of AP2 binding the YXXΦ motif of PBF.

Acidic residues upstream of a dileucine motif have previously been described to be important for endocytosis due to a structure favourable for internalisation and a role in endocytic vesicle formation (40). A discrete putative dileucine motif (562**LL**563) has recently been implicated in basolateral targeting of NIS. Koumarianou and colleagues discovered that the dileucine LL562/563 motif in the C-terminus of NIS was critical to interaction with the AP1μ1A subunit, as part of the polarised trafficking of NIS to the basolateral PM (19). In addition, residues E578 and L583 have been shown to constitute a conserved monoleucine- based sorting motif essential for NIS transport to the basolateral plasma membrane (19,20). Our finding that manipulation of the same residues results in NIS which is retained intracellularly in non-polarised cells therefore supports the observation that dileucine and diacidic motifs are critical to the movement and targeting of NIS in epithelial cells. In this study NanoBiT assays demonstrated the AP2σ2 subunit binds less effectively to both the dileucine and diacidic mutants than WT, revealing an impact of the 2 NIS motifs on interaction with AP2 as well as AP1

Dual redifferentiation therapies, such as those based on combining BRAF and MEK inhibitors, are beginning to show promise in clinical trials (41), but reported mechanisms of resistance to MAPK inhibitors are common (42). Here, our approach was to identify new drug strategies to boost the efficacy of radioiodide therapy based on a greater mechanistic understanding of targetable steps of NIS processing outside of canonical signalling pathways. In particular, CQ is a 4-aminoquinoline which has been used for over 70 years as an antimalarial agent, accumulating preferentially in lysosomes. As such, CQ has been shown to impact multiple cellular processes including autophagy, endo-lysosomal degradation and endocytosis. The relatively rapid (∼8 hour) impact of CQ on RAI uptake suggested that its influence on NIS function may be predominantly via inhibiting endocytic processes. Currently, there are no known drugs capable of altering NIS endocytosis in vivo as ‘experimental’ compounds such as Dynasore are not clinically applicable. Our finding using NanoBRET and cell surface biotinylation assays that CQ inhibited endocytosis to retain NIS at the PM is therefore of significant translatable potential.

A central mechanism, however, underlying radioiodide-refractoriness in thyroid cancer is decreased levels of NIS expression (43), in addition to reduced NIS localisation at the PM. SAHA is a well-characterised FDA-approved HDAC inhibitor (HDACi), induces robust NIS mRNA expression in thyroid cells (44,45), and was shown to improve radioiodide uptake in one of three patients with thyroid cancer in a phase 1 trial (46). For our animal models we wanted to emulate a possible clinical scenario in patients with thyroid cancer: we hypothesized that SAHA treatment would induce NIS expression, and that increased NIS protein might then benefit from endocytosis inhibition to enhance radioiodide uptake. Given that the thyroidal uptake of ^99m^Tc was maximally stimulated in BALB/c mice treated with SAHA and CQ, clinical trials to address whether patients receiving this drug combination at the time of radioiodide therapy uptake more ^131^I would now be timely. Further work is also warranted to determine whether the mechanistic impact of combining CQ with SAHA to potentiate NIS function may additionally occur via blocking the endocytic activity of HDACi-induced genes, given our data that SAHA induced AP2A1 and PICALM expression.

An important clinical observation in our study was the striking correlation between recurrence and the magnitude of endocytic gene dysregulation, including AP2α genes, particularly in patients who received radioiodide treatment. Altering expression of a wide range of endocytic genes is well-recognised to have substantive effects on the maturation and dynamics of clathrin-coated vesicles (47). We thus propose that the extensive dysregulation of endocytic genes discovered in PTC results in NIS mislocalisation away from the PM and reduced radioiodide uptake, leading to a greater number of treatment-resistant tumour cells and increased risk of recurrence. One important finding in support of this was the lack of differences in clinical staging attributes for RAI treatment groups stratified for AP2 genes, suggesting that endocytic genes have no independent impact on tumour pathogenesis. Of particular significance, we further identified a 30-endocytic gene risk score classifier for recurrence, which yielded higher specificity and sensitivity than single gene biomarkers, as well as being an independent predictive factor for recurrence. Based on these findings, we envisage that the 30 endocytic gene classifier represents a promising biomarker for predicting thyroid cancer recurrence, especially in aggressive thyroid cancers requiring radioiodide.

In summary, we delineate endocytic pathways which govern NIS function in thyroid cancer cells. Bioinformatic analyses further revealed extensive dysregulation of endocytic genes in PTC. Although the exact order of events is challenging to discern experimentally, we identify that the overall endocytic process is druggable both in vitro and in vivo. As FDA-approved drugs enhance radionuclide accumulation in the thyroid at realistic therapeutic doses and timepoints, our results indicate that systemic modulation of NIS activity may now be possible in patients. This study offers a new therapeutic approach for RAIR-TC treatment as well as augmenting NIS function for developing radioiodide-based therapies across a broader disease spectrum, including breast cancer.

## ACKNOWLEDGEMENTS

This work was supported by the Department of Defense (BC201532P1 to M.J. Campbell and C.J. McCabe) and Wellcome Trust (RG_05-052 to A. Fletcher). C.J. McCabe also received funding from the Medical Research Council and British Thyroid Association. We acknowledge the labs of K. Pfleger and N. Lambert for kindly providing Venus-tagged constructs, J.E. Blower for in vivo expertise, and D.P. Larner and R.L. Hoare for technical expertise. We further acknowledge support from the Wellcome Trust and EPSRC funded Centre for Medical Engineering at King’s College London (203148/Z/16/Z), the Wellcome Multiuser Equipment Radioanalytical Facility (212885/Z/18/Z), and the EPSRC programme for Next Generation Molecular Imaging and Therapy with Radionuclides (EP/S019901/1).

## Notes

### Competing Interest Statement

The authors have declared no competing interest.

