## Supplementary Information for "Dissecting endocytic mechanisms reveals new molecular targets to enhance sodium iodide symporter activity with clinical relevance to radioiodide therapy"

**This PDF file includes:**

Supplementary Materials and Methods

Figures S1 to S8

### Supplementary Materials and Methods

#### Key resources

| REAGENT | SOURCE | IDENTIFIER |
|---|---|---|
| **Antibodies** | | |
| Rabbit polyclonal anti-AP2A1 | Antibodies.com | Cat# A38279 |
| Rabbit polyclonal anti-AP2M1 | Novus Biologicals | Cat# NBP1-32272 |
| Rabbit polyclonal anti-NIS | Proteintech | Cat# 24324-1-AP |
| Rabbit monoclonal anti-PICALM | Cell Signaling Technology | Cat# 26765 |
| Rabbit polyclonal anti-Na,K-ATPase | Cell Signaling Technology | Cat# 3010 |
| Mouse monoclonal anti-HA | BioLegend | Cat# 901502 |
| Mouse monoclonal anti-β-actin | Sigma-Aldrich | Cat# A1978 |
| **Oligonucleotides** | | |
| AAK1 TaqMan® Gene Expression Assay (human) | ThermoFisher Scientific | Hs00208618_m1 |
| AP2A1 TaqMan® Gene Expression Assay (human) | ThermoFisher Scientific | Hs00900330_m1 |
| AP2M1 TaqMan® Gene Expression Assay (human) | ThermoFisher Scientific | Hs01037584_m1 |
| AP2S1 TaqMan® Gene Expression Assay (human) | ThermoFisher Scientific | Hs00900330_m1 |
| PICALM TaqMan® Gene Expression Assay (human) | ThermoFisher Scientific | Hs00200318_m1 |
| PPIA TaqMan® Gene Expression Assay (human) | ThermoFisher Scientific | Hs04194521_s1 |
| SLC5A5 TaqMan® Gene Expression Assay (human) | ThermoFisher Scientific | Hs00950358_m1 |
| TSHR TaqMan® Gene Expression Assay (human) | ThermoFisher Scientific | Hs01053846_m1 |
| 18S TaqMan® Gene Expression Assay (human) | ThermoFisher Scientific | Hs03003631_g1 |
| ACTB TaqMan® Gene Expression Assay (mouse) | ThermoFisher Scientific | Mm01205647_g1 |
| AP2A1 TaqMan® Gene Expression Assay (mouse) | ThermoFisher Scientific | Mm00475919_m1 |
| AP2M1 TaqMan® Gene Expression Assay (mouse) | ThermoFisher Scientific | Mm05884025_g1 |
| HPRT TaqMan® Gene Expression Assay (mouse) | ThermoFisher Scientific | Mm00446968_m1 |
| NKX2-1 TaqMan® Gene Expression Assay (mouse) | ThermoFisher Scientific | Mm07296387_g1 |
| PAX8 TaqMan® Gene Expression Assay (mouse) | ThermoFisher Scientific | Mm00440623_m1 |
| PICALM TaqMan® Gene Expression Assay (mouse) | ThermoFisher Scientific | Mm00525455_m1 |
| PTTG1IP TaqMan® Gene Expression Assay (mouse) | ThermoFisher Scientific | Mm00521473_m1 |
| SLC5A5 TaqMan® Gene Expression Assay (mouse) | ThermoFisher Scientific | Mm01351811_m1 |
| TSHR TaqMan® Gene Expression Assay (mouse) | ThermoFisher Scientific | Mm00442027_m1 |
| ON-TARGETplus Human AAK1 siRNA | Horizon Discovery | Cat# L-005300-02-0005 |
| ON-TARGETplus Human AP2A1 siRNA | Horizon Discovery | Cat# L-012492-00-0005 |
| ON-TARGETplus Human AP2A2 siRNA | Horizon Discovery | Cat# L-012812-00-0005 |
| ON-TARGETplus Human AP2M1 siRNA | Horizon Discovery | Cat# L-008170-00-0005 |
| ON-TARGETplus Human AP2S1 siRNA | Horizon Discovery | Cat# L-011833-01-0005 |
| ON-TARGETplus Non-targeting Pool siRNA | Horizon Discovery | Cat# D-001810-10-05 |
| LgBiT (PCR primers; N-terminal)<br>5’-GCCAAGCTTACCATGGTCTTCACACTCGAA-3’<br>5’-GGCGGTACCACTGTTGATGGTTACTCG-3’ | Sigma-Aldrich | Custom synthesis |
| LgBiT (PCR primers; C-terminal)<br>5’-GCCTCTAGAGTCTTCACACTCGAAGAT-3’<br>5’-GGCGGCGGGCCCTTAGCTGTTGATGGTTAC-3’ | Sigma-Aldrich | Custom synthesis |
| AP2S1 (PCR primers; N-terminal tag with LgBiT)<br>5’-GCCGGTACCATCCGCTTTATCCTCATC-3’<br>5’-GCCTCTAGATCACTCCAGGGACTGTAG-3’ | Sigma-Aldrich | Custom synthesis |
| AP2S1 (PCR primers; C-terminal tag with LgBiT)<br>5’-GCCGGTACCACCATGATCCGCTTTATCCTC-3’<br>5’-GCCTCTAGACTCCAGGGACTGTAGCAT-3’ | Sigma-Aldrich | Custom synthesis |

| | | |
|---|---|---|
| AP2S1 (PCR primers; N-terminal tag with SmBiT)<br>5’-GCCGGTACCACCATGGTGACCGGCTACCGGCTGTTCGAGGAGATTCTCATCCGCTTTATCCTCA-3’<br>5’-GCCGCGCCGCCGCCTCTAGATCACTCCAGGGACTGT-3’ | Sigma-Aldrich | Custom synthesis |
| AP2S1 (PCR primers; C-terminal tag with SmBiT)<br>5’-GCCGCCGCCGCCGGTACCACCATGATCCGCTTTATCC-3’<br>5’-GCCTCTAGATTACAGAATCTCCTCGAACAGCCGGTAGCCGGTCACCTCCAGGGACTGTAGC-3’ | Sigma-Aldrich | Custom synthesis |
| PBF (PCR primers; N-terminal tag with LgBiT)<br>5’- GCCGGTACCGCGCCCGGAGTGGCCCG-3’<br>5’- GCCTCTAGATTAGTTGTTTTCAAATCT-3’ | Sigma-Aldrich | Custom synthesis |
| PBF (PCR primers; C-terminal tag with LgBiT)<br>5’-GCCGGTACCACCATGGCGCCCGGAGTGGC-3’<br>5’-GCCTCTAGAGTTGTTTTCAAATCTAGC-3’ | Sigma-Aldrich | Custom synthesis |
| PBF (PCR primers; N-terminal tag with SmBiT)<br>5’-GCCGGTACCACCATGGTGACCGGCTACCGGCTGTTCGAGGAGATTCTCGCGCCCGGAGTGGCCC-3’<br>5’-GCCGCGCCGCCGCCTCTAGATTAGTTGTTTTCAAATC-3’ | Sigma-Aldrich | Custom synthesis |
| PBF (PCR primers; C-terminal tag with SmBiT)<br>5’-GCCGCCGCCGCCGGTACCACCATGGCGCCCGGAGTG-3’<br>5’-GCCTCTAGAGAGAATCTCCTCGAACAGCCGGTAGCCGGTCACGTTGTTTTCAAATCTAG-3’ | Sigma-Aldrich | Custom synthesis |
| NIS (mutagenesis primers; L562A and L563A)<br>5’-CCCCGGGAGCGGCGTGGTGGGAC-3’<br>5’-GTCCCACCACGCCGCTCCCGGGG-3’ | Sigma-Aldrich | Custom synthesis |
| NIS (mutagenesis primers; E578A and E579A)<br>5’-GGATGGCCACTGCTGCCTTGGGG-3’<br>5’-CCCCAAGGCAGCAGTGGCCATCC-3’ | Sigma-Aldrich | Custom synthesis |
| AP2S2 (mutagenesis primers; V88D)<br>5’-AGGCCATTCACAACTTCGACGAGGTCTTAAACGAATATTT-3’<br>5’-AAATATTCGTTTAAGACCTCGTCGAAGTTGTGAATGGCCT-3’ | Sigma-Aldrich | Custom synthesis |
| AP2S2 (mutagenesis primers; L103D)<br>5’-CAATGTCTGTGAACTGGACAGCGTGTTCAACTTCTACAAG-3’<br>5’-CTTGTAGAAGTTGAACACGCTGTCCAGTTCACAGACATTG-3’ | Sigma-Aldrich | Custom synthesis |
| **Recombinant DNA** | | |
| pcDNA3.1-NIS-HA | Smith VE et al., 2009 | Supp Ref (1) |
| pcDNA3.1-PBF-HA | Smith VE et al., 2009 | Supp Ref (1) |
| pcDNA3.1-NIS-SmBiT | Read ML et al., 2022 | Supp Ref (2) |
| pcDNA3.1-NIS-LgBiT | This paper | N/A |
| pcDNA3.1-PBF-SmBiT | This paper | N/A |
| pcDNA3.1-LgBiT (N-terminal; HindIII/KpnI) | This paper | N/A |
| pcDNA3.1-LgBiT (C-terminal; XbaI/ApaI) | This paper | N/A |
| pcDNA3.1-AP2S1-LgBiT | This paper | N/A |
| pcDNA3.1-LgBiT-AP2S1 | This paper | N/A |
| pcDNA3.1-AP2S1-SmBiT | This paper | N/A |
| pcDNA3.1-SmBiT-AP2S1 | This paper | N/A |
| pcDNA3.1-NIS(L562A/L563A)-SmBiT | This paper | N/A |
| pcDNA3.1-NIS(E578A/E579A)-SmBiT | This paper | N/A |
| pcDNA3.1-LgBiT-AP2S1(V88D) | This paper | N/A |
| pcDNA3.1-LgBiT-AP2S1(L103S) | This paper | N/A |
| pcDNA3.1-NIS(L562A/L563A)-HA | This paper | N/A |

| | | |
|---|---|---|
| pcDNA3.1-NIS(E578A/E579A)-HA | This paper | N/A |
| pcDNA3.1-AP2S1(V88D) | This paper | N/A |
| pcDNA3.1-AP2S1(L103S) | This paper | N/A |
| pcDNA3.1-Rab1-Venus | Kevin Pfleger’s lab | N/A |
| pcDNA3.1-Rab4-Venus | Kevin Pfleger’s lab | N/A |
| pcDNA3.1-Rab6-Venus | Kevin Pfleger’s lab | N/A |
| pcDNA3.1-Rab8-Venus | Kevin Pfleger’s lab | N/A |
| pcDNA3.1-Rab9-Venus | Kevin Pfleger’s lab | N/A |
| pcDNA3.1-Kras-Venus | Nevin Lambert’s lab | N/A |
| pcDNA3.1-Rab5-Venus | Nevin Lambert’s lab | N/A |
| pcDNA3.1-Rab7-Venus | Nevin Lambert’s lab | N/A |
| pcDNA3.1-Rab11-Venus | Nevin Lambert’s lab | N/A |
| pcDNA3.1-NIS-Nluc | ThermoFisher Scientific | N/A |
| pCMV3-AP2S1-untagged | Sino Biological Europe GmbH | Cat# HG12478-UT |

#### Construction of NanoBiT LgBiT- and SmBiT-tagged genes

The LgBiT tag was amplified with Q5 DNA Polymerase (NEB) using either N- or C-terminal LgBiT primers (see Key resources; NanoBiT PPI starter system; N2014). Following gel extraction (QIAquick kit; Qiagen), PCR products were digested with HindIII and KpnI (Promega) for N-terminal tagging or with XbaI and ApaI (Promega) for C-terminal tagging, prior to ligation to pcDNA3.1(+) (ThermoFisher Scientific) linearised with the same pairs of restriction enzymes. Next, AP2S1 cDNA was amplified using specific primers for N- and C-terminal LgBiT tagging (see Key resources). PCR products were digested with KpnI and XbaI, and then ligated (T4 DNA ligase; NEB) into LgBiT-containing vectors [pcDNA3.1-LgBiT (N-terminal); pcDNA3.1-LgBiT (C-terminal)] linearised with the same restriction enzymes. AP2S1 and PBF cDNA were amplified with the SmBiT tag directly using appropriate PCR primers (see Key resources) and ligated into the empty pcDNA3.1(+) vector linearised with the same restriction enzymes.

#### TCGA and GEO datasets

RNA-seq data for 59 normal thyroid and 501 PTC TCGA samples were analyzed (Broad GDAC Firehose, doi:10.7908/C11G0KM9). Normalised gene expression values were transformed as $X=\log2(X+1)$ where X represents the normalized fragments per kilobase transcript per million mapped reads (FPKM) values. Differential gene expression analysis was also performed using the GEO2R interactive web tool in GEO (3) to investigate endocytic genes in thyroid cancer, including analysis of the GEO dataset GSE60542 (4). Heatmaps were constructed using Morpheus (Broad Institute; https://software.broadinstitute.org/morpheus). Functional gene classification of TCGA RNA-seq data was performed using DAVID (5,6) and ToppGene (7).

#### Statistical analyses

All results were obtained from triplicate biological experiments unless otherwise indicated. For comparison between two groups, data were subjected to the Student’s t-test, and for multiple comparisons one-way ANOVA was used with either Dunnett's or Tukey's post-hoc test. Kruskal-Wallis and Spearman's correlation tests were performed on non-parametric data. P-values were adjusted using the Benjamini-Hochberg FDR correction procedure to correct for multiple comparisons. Dunn's multiple comparison post-hoc testing was used after Kruskal-Wallis tests to determine significance between datasets in groups of 3 or more. Fisher's exact test was used to determine the significance of nonrandom associations between two categorical variables. $P < 0.05$ was considered significant. All *P*-values reported from statistical tests were two-sided.

**A**

#### TPC-1

**Laboratory Report**

| | |
|---|---|
| Test Requested | Cell Line Authentication |
| Case Number | C-25115d |
| Date Sample Received | 08/04/2022 |
| Date Sample Tested | 11/04/2022 |
| Date Sample Reported | 19/04/2022 |

| Sample Name | Sample/Comparison Profile Source | Sample Number | DNA Number |
|---|---|---|---|
| TPC-1 | University of Birmingham | S-1057716 | D-1057716 |
| TPC-1 | Cellosaurus Database | N/A | N/A |

**Table of Allelic Data**

| | Genotypes | | |
|---|---|---|---|
| STR Locus | TPC-1 (Test Sample) | TPC-1 (Comparison Sample) | Match vs. Mis-Match |
| D5 | 8 \| 10 | 8 \| 10 | Match |
| D13 | 11 \| 12 | 11 \| 12 | Match |
| D7 | 11 \| 11 | 11 \| 11 | Match |
| D16 | 9 \| 9 | 9 \| 9 | Match |
| vWA | 14 \| 18 | 14 \| 18 | Match |
| Amel | X \| X | X \| X | Match |
| TPOX | 11 \| 11 | 11 \| 11 | Match |
| CSF1PO | 11 \| 12 | 11 \| 12 | Match |
| THO1 | 9 \| 9 | 9 \| 9 | Match |

Matching Percentage: **100%**
Outcome: **Related**

**B**

## 8505C

**Laboratory Report**

| | |
|---|---|
| Test Requested | Cell Line Authentication |
| Case Number | C-25115e |
| Date Sample Received | 08/04/2022 |
| Date Sample Tested | 11/04/2022 |
| Date Sample Reported | 19/04/2022 |

| Sample Name | Sample/Comparison Profile Source | Sample Number | DNA Number |
|---|---|---|---|
| 8505C | University of Birmingham | S-1057717 | D-1057717 |
| 8505C | DSMZ Database | N/A | N/A |

**Table of Allelic Data**

| | Genotypes | | |
|---|---|---|---|
| STR Locus | 8505C (Test Sample) | 8505C (Comparison Sample) | Match vs. Mis-Match |
| D5 | 10 \| 11 | 10 \| 11 | Match |
| D13 | 13 \| 13 | 13 \| 13 | Match |
| D7 | 10 \| 10 | 10 \| 10 | Match |
| D16 | 12 \| 12 | 12 \| 12 | Match |
| vWA | 17 \| 19 | 17 \| 19 | Match |
| Amel | X \| X | X \| X | Match |
| TPOX | 10 \| 11 | 10 \| 11 | Match |
| CSF1PO | 12 \| 13 | 12 \| 13 | Match |
| THO1 | 6 \| 9 | 6 \| 9 | Match |

Matching Percentage: **100%**
Outcome: **Related**

**C**

#### HeLa

**Laboratory Report**

| | |
|---|---|
| Test Requested | Cell Line Authentication |
| Case Number | C-25115f |
| Date Sample Received | 08/04/2022 |
| Date Sample Tested | 11/04/2022 |
| Date Sample Reported | 19/04/2022 |

| Sample Name | Sample/Comparison Profile Source | Sample Number | DNA Number |
|---|---|---|---|
| HeLa | University of Birmingham | S-1057718 | D-1057718 |
| HeLa | DSMZ Database | N/A | N/A |

**Table of Allelic Data**

| | Genotypes | | |
|---|---|---|---|
| STR Locus | HeLa (Test Sample) | HeLa (Comparison Sample) | Match vs. Mis-Match |
| D5 | 11 \| 12 | 11 \| 12 | Match |
| D13 | 12 \| 13.3 | 12 \| 13.3 | Match |
| D7 | 8 \| 12 | 8 \| 12 | Match |
| D16 | 9 \| 10 | 9 \| 10 | Match |
| vWA | 16 \| 18 | 16 \| 18 | Match |
| Amel | X \| X | X \| X | Match |
| TPOX | 8 \| 12 | 8 \| 12 | Match |
| CSF1PO | 9 \| 10 | 9 \| 10 | Match |
| THO1 | 7 \| 7 | 7 \| 7 | Match |

Matching Percentage: **100%**
Outcome: **Related**

**D**

#### HEK293

**Laboratory Report**

| | |
|---|---|
| Test Requested | Cell Line Authentication |
| Case Number | C-25115h |
| Date Sample Received | 08/04/2022 |
| Date Sample Tested | 11/04/2022 |
| Date Sample Reported | 19/04/2022 |

| Sample Name | Sample/Comparison Profile Source | Sample Number | DNA Number |
|---|---|---|---|
| HEK293 | University of Birmingham | S-1057720 | D-1057720 |
| HEK-293.2sus | DSMZ Database | N/A | N/A |

**Table of Allelic Data**

| | Genotypes | | |
|---|---|---|---|
| STR Locus | HEK293 (Test Sample) | HEK-293.2sus (Comparison Sample) | Match vs. Mis-Match |
| D5 | 8 \| 8 | 8 \| 8 | Match |
| D13 | 12 \| 14 | 12 \| 14 | Match |
| D7 | 11 \| 11 | 11 \| 12 | Mis-Match |
| D16 | 9 \| 13 | 9 \| 13 | Match |
| vWA | 16 \| 19 | 16 \| 19 | Match |
| Amel | X \| X | X \| X | Match |
| TPOX | 11 \| 11 | 11 \| 11 | Match |
| CSF1PO | 12 \| 12 | 12 \| 12 | Match |
| THO1 | 7 \| 9.3 | 7 \| 9.3 | Match |

Matching Percentage: **94%**
Outcome: **Related**

**Figure S1.** Representative STR profiles of cell lines used in study.

Examples of cell line authentication reports for **A,** TPC-1, **B,** 8505C, **C,** HeLa and **D,** HEK293 cells generated by NorthGene (Biofortuna). DNA profiles were compared to profiles located on either the Cellosaurus or DSMZ database as indicated. Analysis was conducted using 8 loci allowing identical matches an approximate power of discrimination of 1 in 1,000,000,000. Cell lines were considered to be related with a matching percentage > 80%.

A

| Gene | BRAF-like, RAI (n = 137, FDR < 12%) | | BRAF-like, Non-RAI (n = 94) | |
|---|---|---|---|---|
| | p | $N_{REC}$ | p | $N_{REC}$ |
| *ACTB* | 0.593 | 12,9 | 0.985 | 1,1 |
| *ACTR3* | 0.804 | 11,10 | 0.146 | 2,0 |
| *AP2A2* | 0.043 | 15,6 | 0.956 | 1,1 |
| *AP2B1* | 0.295 | 13,8 | 0.969 | 1,1 |
| *ARF6* | 0.031 | 15,6 | 0.146 | 2,0 |
| *ARPC4* | 0.803 | 11,10 | 0.969 | 1,1 |
| *CTTN* | 0.032 | 15,6 | 0.146 | 2,0 |
| *EHD1* | 0.929 | 11,10 | 0.981 | 1,1 |
| *HLA-A* | 0.009 | 16,5 | 0.998 | 1,1 |
| *PDCD6IP* | 0.871 | 10,11 | 0.905 | 2,0 |
| *PICALM* | 0.432 | 9,12 | 0.134 | 2,0 |
| *RAB5C* | 0.002 | 17,4 | 0.956 | 1,1 |
| *SRC* | 0.126 | 14,7 | 0.905 | 1,1 |
| *TF* | 0.228 | 8,13 | 0.969 | 1,1 |

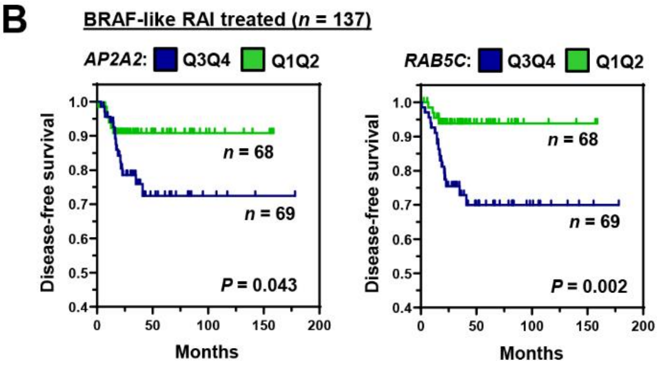


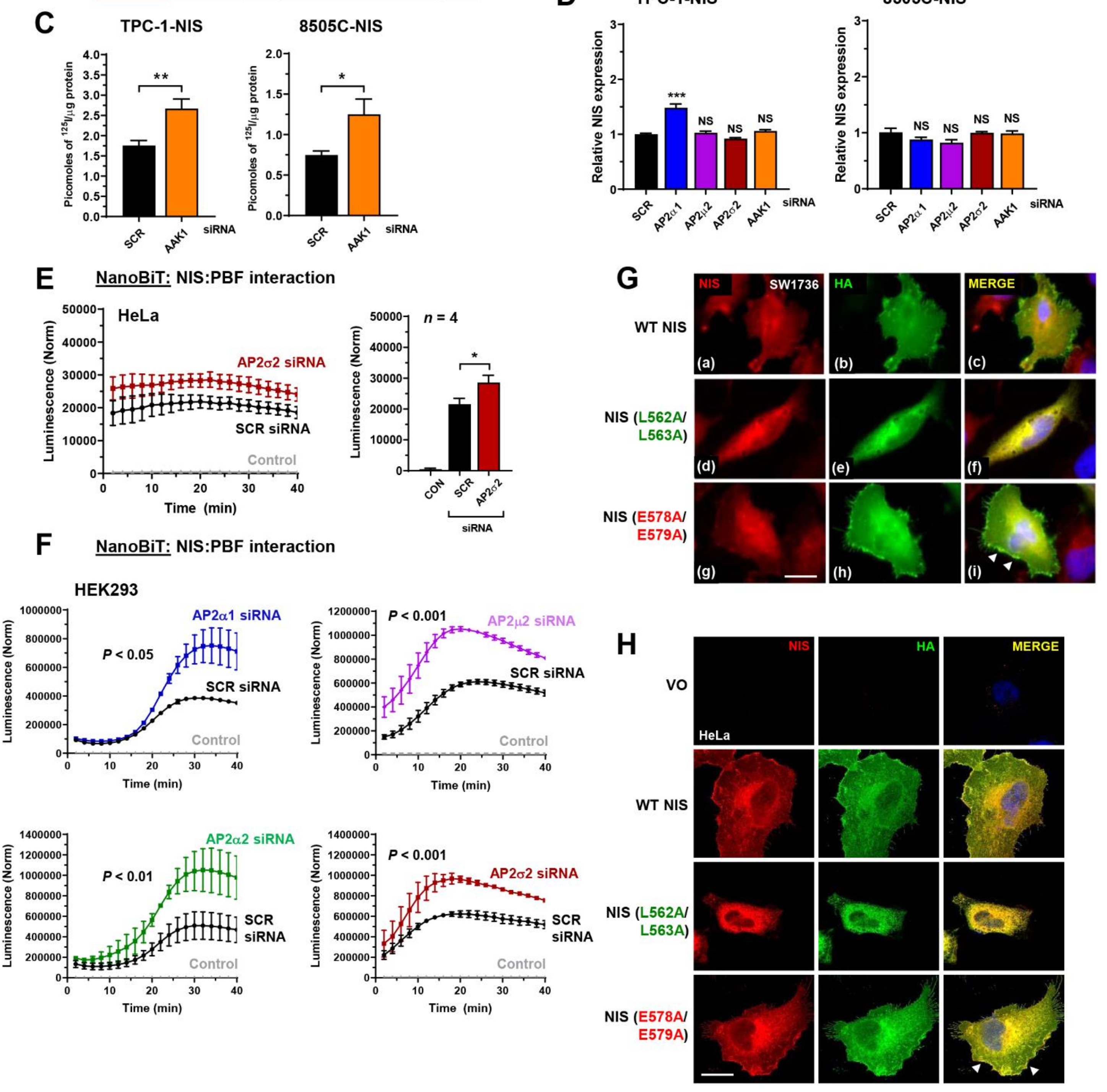

**Figure S2.** Association of NIS interactors with recurrence.

**A,** Kaplan-Meier analysis of the RAI-treated (*n* = 137) and non-RAI treated (*n* = 94) THCA cohorts stratified on high (Q3Q4) versus low (Q1Q2) tumoral expression; log-rank test. Significance indicated by *P*-values with FDR < 12%. $N_{REC}$ = number of recurrent cases in cohort with either high (left) or low (right) tumoral expression. Green = $P < 0.05$; Yellow = highest percentage of stratified recurrent cases.

**B,** Representative Kaplan-Meier analysis of DFS for the BRAF-like, RAI-treated THCA cohort stratified on high (Q3Q4) versus low (Q1Q2) tumoral expression of indicated endocytosis genes; log-rank test. Number (*n*) of patients per expression sub-group (high/low) and *P*-values are shown.

**C,** RAI uptake of TPC-1-NIS and 8505C-NIS cells transfected with AAK1 versus scrambled (scr) siRNA.

**D,** Relative mRNA levels of NIS in TPC-1-NIS and 8505C-NIS cells transfected with siRNA specific for indicated AP2 genes and AAK1.

**E,** NanoBiT evaluation of protein: protein interaction between NIS and PBF in living HeLa cells transfected with AP2$\sigma$2 siRNA. (*right*) Normalised NanoBiT assay results at 20 minutes post-addition of Nano-Glo live cell assay substrate (*n* = 4).

**F,** Same as **E** but HEK293 cells transfected with AP2$\alpha$1, AP2$\alpha$2, AP2$\mu$2 and AP2$\sigma$2 siRNA (*n* = 3 - 4).

**G** and **H,** Confocal imaging in SW1736 (**G**) and HeLa (**H**) cells transfected with wild-type (WT) NIS, NIS mutant L562/L563A or NIS mutant E578A/E579A. Confocal images represent NIS expression (red), HA expression (green) and a merged image (yellow). Arrows (white) highlight PM regions with greater NIS localisation. Scale bar – 20 μm.

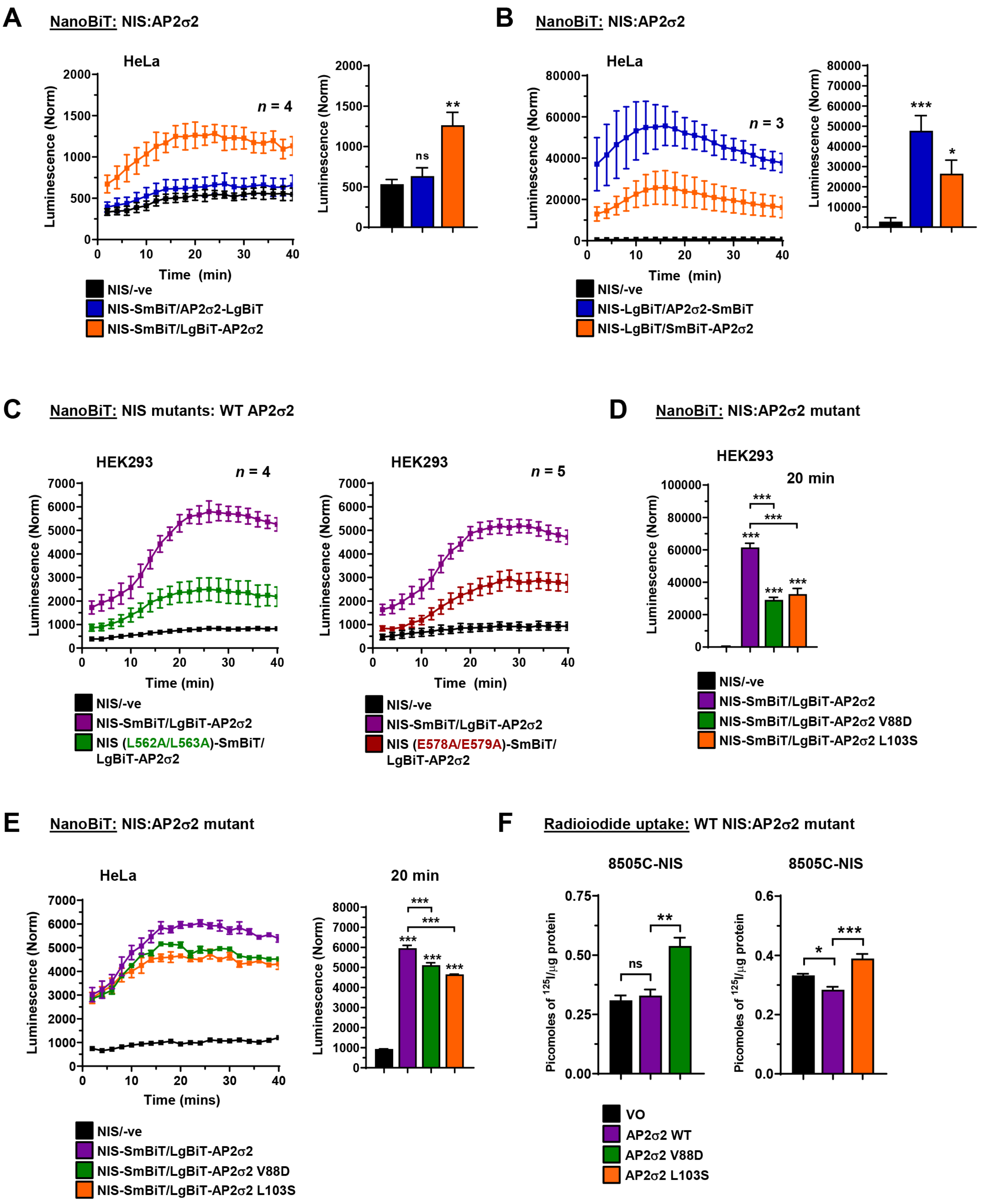

A NanoBiT: NIS:AP2σ2
HeLa
n = 4
Luminescence (Norm)
Time (min)
ns
**
NIS/-ve
NIS-SmBiT/AP2σ2-LgBiT
NIS-SmBiT/LgBiT-AP2σ2
B NanoBiT: NIS:AP2σ2
HeLa
n = 3
***
*
NIS/-ve
NIS-LgBiT/AP2σ2-SmBiT
NIS-LgBiT/SmBiT-AP2σ2
C NanoBiT: NIS mutants: WT AP2σ2
HEK293
n = 4
n = 5
NIS/-ve
NIS-SmBiT/LgBiT-AP2σ2
NIS (L562A/L563A)-SmBiT/ LgBiT-AP2σ2
NIS (E578A/E579A)-SmBiT/ LgBiT-AP2σ2
D NanoBiT: NIS:AP2σ2 mutant
HEK293
20 min
NIS-SmBiT/LgBiT-AP2σ2 V88D
NIS-SmBiT/LgBiT-AP2σ2 L103S
E NanoBiT: NIS:AP2σ2 mutant
HeLa
Time (mins)
20 min
F Radioiodide uptake: WT NIS:AP2σ2 mutant
8505C-NIS
Picomoles of 125I/μg protein
VO
AP2σ2 WT
AP2σ2 V88D
AP2σ2 L103S

**Figure S3.** Kinetic profiling of the interaction between NIS and AP2σ2 in live cells.

**A,** Live cell kinetic measurement using the NanoBiT assay to evaluate protein-protein interactions between NIS and AP2σ2 tagged with LgBiT in HeLa cells. (*right*) NanoBiT assay results at 20 minutes post-addition of Nano-Glo substrate.

**B,** Same as **A** but AP2σ2 tagged with SmBiT.

**C,** Same as **A** but using NIS mutants L562A/L563A (*left*) and E578A/E579A (*right*) in HEK293 cells.

**D,** Same as **A** but using AP2σ2 mutants V88D and L103S in HEK293 cells. NanoBiT assay results are shown at 20 minutes post-addition of Nano-Glo substrate.

**E,** Same as **D** but in HeLa cells.

**F,** RAI uptake in 8505C-NIS cells transfected with WT AP2σ2, AP2σ2 mutant V88D and AP2σ2 mutant L103S.

Data presented as mean ± S.E.M., $n = 4 - 5$, one-way ANOVA followed by Tukey's post hoc test (ns, not significant; *$P < 0.05$; **$P < 0.01$; ***$P < 0.001$).

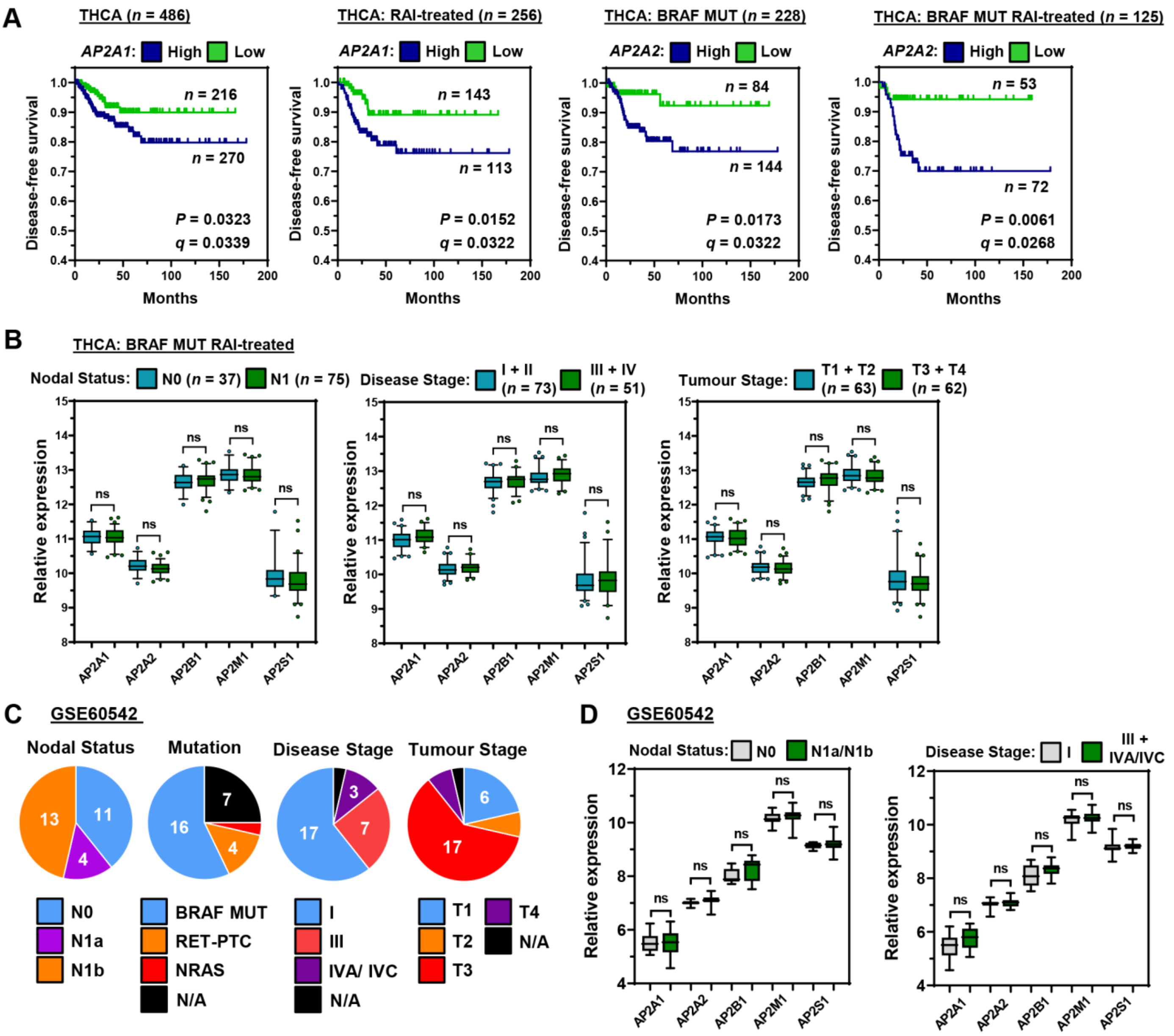


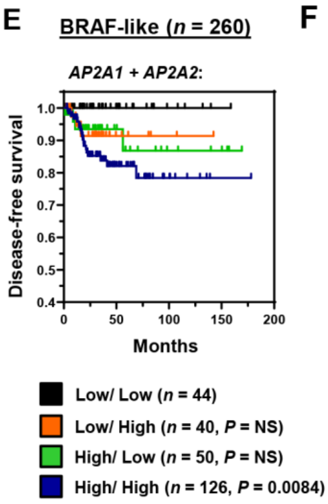


**F** THCA: BRAF-like RAI-treated (*n* = 137)

| Gene #1 | Gene #2 | Expression Group | | | | Disease-Free Survival (L/L vs H/H) | | Frequency of Recurrence (L/L vs H/H) | |
|---|---|---|---|---|---|---|---|---|---|
| | | Low/Low | | High/High | | | | | |
| | | $N_{tot}$ | $N_{rec}$ | $N_{tot}$ | $N_{rec}$ | *P* | *q* | *P* | *q* |
| *AP2A1* | *AP2A2* | 29 | 0 | 62 | 15 | 0.006 | 0.038 | 0.002 | 0.013 |
| *AP2A1* | *AP2B1* | 31 | 1 | 51 | 11 | 0.034 | 0.071 | 0.026 | 0.055 |
| *AP2A1* | *AP2M1* | 34 | 2 | 42 | 5 | 0.411 | 0.432 | 0.450 | 0.473 |
| *AP2A1* | *AP2S1* | 23 | 2 | 55 | 8 | 0.619 | 0.557 | 0.714 | 0.643 |
| *AP2A2* | *AP2B1* | 31 | 2 | 39 | 12 | 0.020 | 0.063 | 0.015 | 0.047 |
| *AP2A2* | *AP2M1* | 42 | 1 | 38 | 4 | 0.151 | 0.238 | 0.185 | 0.291 |
| *AP2A2* | *AP2S1* | 32 | 2 | 54 | 8 | 0.260 | 0.328 | 0.310 | 0.391 |

**Figure S4.** Lack of association between AP2 genes and cancer staging attributes.

**A,** Representative Kaplan-Meier analysis of DFS for the THCA TCGA cohorts as indicated stratified on high versus low tumoral expression for either *AP2A1* or *AP2A2*; log-rank test. Number ($n$) of patients per expression sub-group (high/low), $P$-values and $q$-values are shown.

**B,** Box and whisker plots showing expression ($\log_2$) of indicated AP2 genes in the BRAF MUT, RAI-treated THCA cohort stratified on nodal status (*left*), disease stage (*middle*) and tumour stage (*right*); $P$-values determined by Mann-Whitney test and adjusted using the Benjamini-Hochberg FDR correction procedure (NS, not significant).

**C,** Pie charts showing the cancer staging and tumour mutational characteristics of patients in the GSE60542 cohort ($n$ = 28).

**D,** Box and whisker plots showing expression ($\log_2$) of indicated AP2 genes in the GSE60542 cohort stratified on nodal status (*left*) and disease stage (*right*); $P$-values determined by Mann-Whitney test and adjusted using the Benjamini-Hochberg FDR correction procedure (NS, not significant).

**E,** Representative Kaplan-Meier analysis of DFS for the BRAF-like THCA cohort stratified on high versus low tumoral expression of both *AP2A1* and *AP2A2*; log-rank test. Number ($n$) of patients per expression sub-group (high/low), $P$-values are shown.

**F,** Kaplan-Meier analysis of the BRAF-like, RAI-treated THCA cohort stratified on high (H/H) versus low (L/L) tumoral expression of either *AP2A1* or *AP2A2* combined with other AP2 genes; log-rank test. Total number ($N_{tot}$) of patients and number of recurrent cases ($N_{rec}$) in each stratified group are shown; significance indicated by $P$- and $q$-values. Fisher's exact test used to determine statistical associations for incidence of recurrence between groups with high (H/H) versus low (L/L) tumoural gene expression.

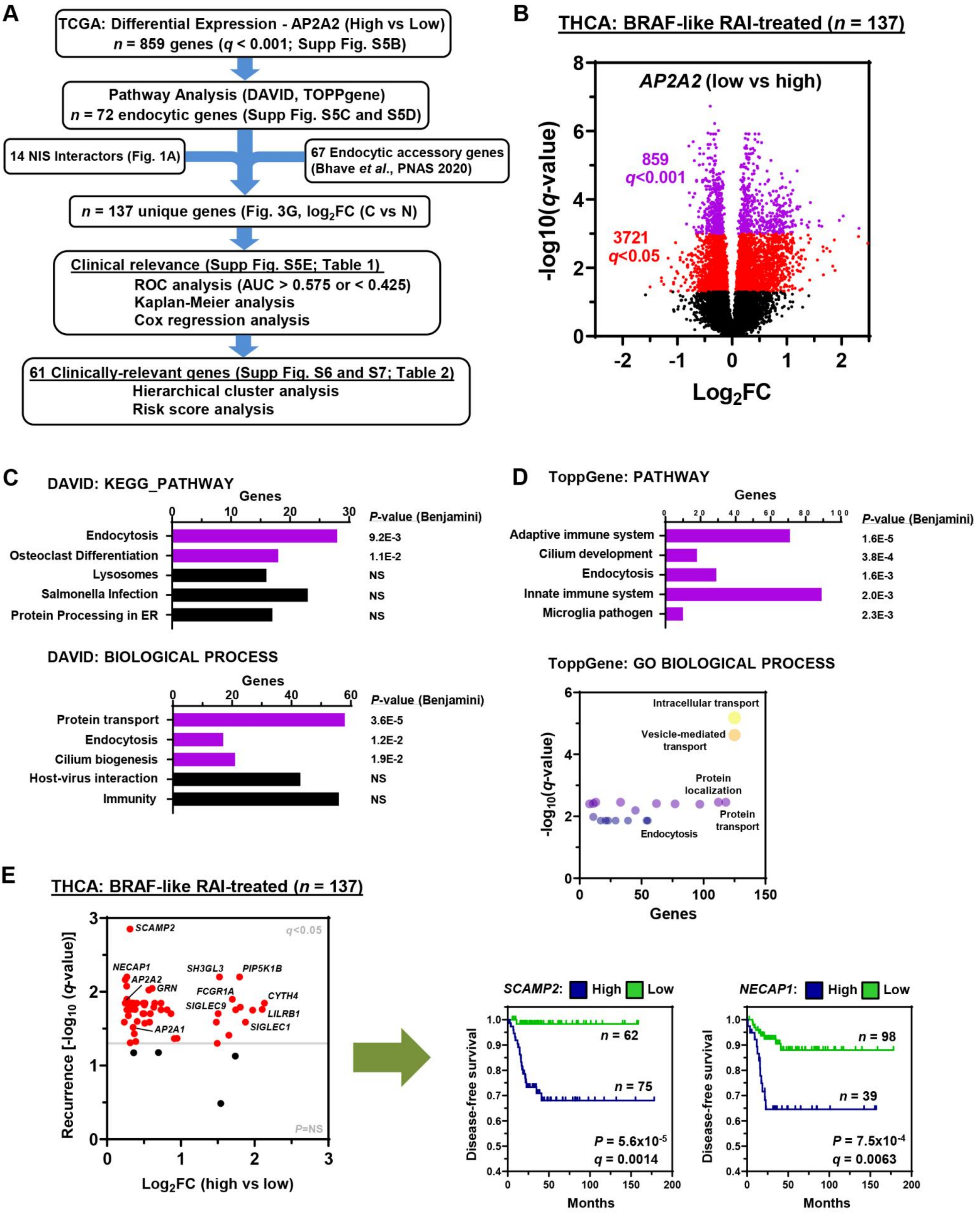
A
TCGA: Differential Expression - AP2A2 (High vs Low)
n = 859 genes (q < 0.001; Supp Fig. S5B)
Pathway Analysis (DAVID, TOPPgene)
n = 72 endocytic genes (Supp Fig. S5C and S5D)
14 NIS Interactors (Fig. 1A)
67 Endocytic accessory genes (Bhave et al., PNAS 2020)
n = 137 unique genes (Fig. 3G, log2FC (C vs N)
Clinical relevance (Supp Fig. S5E; Table 1)
ROC analysis (AUC > 0.575 or < 0.425)
Kaplan-Meier analysis
Cox regression analysis
61 Clinically-relevant genes (Supp Fig. S6 and S7; Table 2)
Hierarchical cluster analysis
Risk score analysis
B
THCA: BRAF-like RAI-treated (n = 137)
AP2A2 (low vs high)
859
q<0.001
3721
q<0.05
-log10(q-value)
Log2FC
C
DAVID: KEGG_PATHWAY
Genes
P-value (Benjamini)
Endocytosis 9.2E-3
Osteoclast Differentiation 1.1E-2
Lysosomes NS
Salmonella Infection NS
Protein Processing in ER NS
DAVID: BIOLOGICAL PROCESS
Protein transport 3.6E-5
Endocytosis 1.2E-2
Cilium biogenesis 1.9E-2
Host-virus interaction NS
Immunity NS
D
ToppGene: PATHWAY
Genes
P-value (Benjamini)
Adaptive immune system 1.6E-5
Cilium development 3.8E-4
Endocytosis 1.6E-3
Innate immune system 2.0E-3
Microglia pathogen 2.3E-3
ToppGene: GO BIOLOGICAL PROCESS
Intracellular transport
Vesicle-mediated transport
Protein localization
Protein transport
Endocytosis
-log10(q-value)
Genes
E
THCA: BRAF-like RAI-treated (n = 137)
SCAMP2
q<0.05
NECAP1
AP2A2
GRN
SH3GL3
PIP5K1B
FCGR1A
SIGLEC9
CYTH4
LILRB1
SIGLEC1
AP2A1
P=NS
Recurrence [-log10 (q-value)]
Log2FC (high vs low)
SCAMP2: High Low
n = 62
n = 75
P = 5.6x10^-5
q = 0.0014
Disease-free survival
Months
NECAP1: High Low
n = 98
n = 39
P = 7.5x10^-4
q = 0.0063

**Figure S5.** Identification of clinical endocytic gene biomarkers for PTC recurrence.

**A,** Summary of bioinformatic pipeline used to identify potential clinical biomarkers of PTC recurrence.

**B,** Volcano plot comparing $\log_2$FC with *q*-value (-log base 10) for the BRAF-like, RAI-treated THCA dataset [high versus low AP2A2 expression; $n = 137$]. Top 859 genes ($q < 0.001$; purple spots) were filtered based on functional classification.

**C,** DAVID functional classification of top 859 differentially expressed genes as described in **A**. KEGG pathway (*upper*) and biological process (*lower*) categories of greatest significance ($P < 0.05$ or lower; purple bars) and number of genes per category are indicated.

**D,** Same as **C** but ToppGene functional classification used to filter the top 859 differentially expressed genes ($q < 0.001$).

**E,** Volcano plot illustrating $\log_2$FC in BRAF-like, RAI-treated THCA cohort (high versus low expression) compared to *q*-value (-log base 10) of recurrence for 61 endocytosis genes ($q < 0.05$). (*right*) Representative Kaplan-Meier analysis of DFS for the BRAF-like, RAI treated THCA cohorts stratified on high versus low tumoural expression of *SCAMP2* and *NECAP1*; log-rank test. Number (*n*) of patients per expression sub-group (high/low), *P*-values and *q*-values are shown.

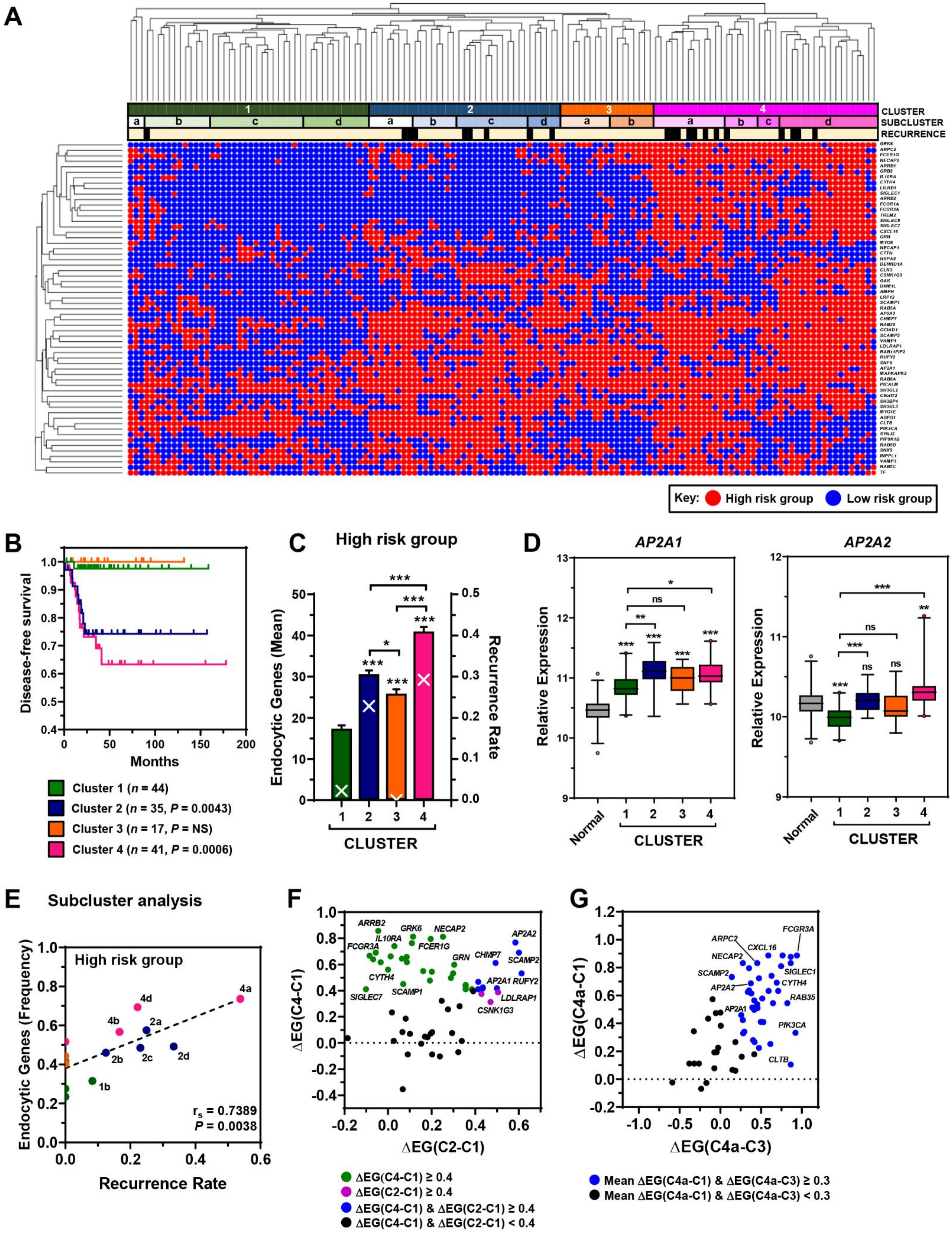
A
CLUSTER
SUBCLUSTER
RECURRENCE
Key: High risk group Low risk group
B
Disease-free survival
Months
Cluster 1 (n = 44)
Cluster 2 (n = 35, P = 0.0043)
Cluster 3 (n = 17, P = NS)
Cluster 4 (n = 41, P = 0.0006)
C
High risk group
Endocytic Genes (Mean)
Recurrence Rate
CLUSTER
D
AP2A1
AP2A2
Relative Expression
Normal
CLUSTER
E
Subcluster analysis
High risk group
Endocytic Genes (Frequency)
Recurrence Rate
rs = 0.7389
P = 0.0038
F
ΔEG(C4-C1)
ΔEG(C2-C1)
ΔEG(C4-C1) ≥ 0.4
ΔEG(C2-C1) ≥ 0.4
ΔEG(C4-C1) & ΔEG(C2-C1) ≥ 0.4
ΔEG(C4-C1) & ΔEG(C2-C1) < 0.4
G
ΔEG(C4a-C1)
ΔEG(C4a-C3)
Mean ΔEG(C4a-C1) & ΔEG(C4a-C3) ≥ 0.3
Mean ΔEG(C4a-C1) & ΔEG(C4a-C3) < 0.3

**Figure S6.** Dysregulated endocytic genes in high-risk group correlate with recurrence rate.

**A,** Hierarchical cluster analysis of the BRAF-like, RAI-treated THCA cohort ($n$ = 137) based on endocytic genes ($n$ = 61) stratified into high and low risk expression groups. Expression cut-off values for stratifying into high and low risk groups determined by ROC analysis (**Table 1B**). Patients were stratified into 4 major clusters (1 to 4) and 14 subclusters (i.e. 1a to 4d). Patients with recurrent disease are indicated (black squares).

**B,** Representative Kaplan-Meier analysis of DFS for the BRAF-like, RAI treated THCA cohort stratified into patient clusters 1 to 4; log-rank test. Number ($n$) of patients per expression sub-group (high/low), and $P$-values are shown.

**C,** Mean number of dysregulated endocytic genes stratified into the high-risk group (bars; left y-axis) and recurrence rate (white crosses; right y-axis) in patient clusters 1 to 4 ($n$ = 17 – 44).

**D,** Box and whisker plots showing expression ($\log_2$) of *AP2A1* (*left*) and *AP2A2* (*right*) in the BRAF-like, RAI-treated THCA cohort stratified into patient clusters 1 to 4 versus normal.

**E,** Correlation analysis between frequency of dysregulated endocytic genes in high-risk group versus recurrence rate in patients stratified into 14 subclusters; Spearman's rank correlation.

**F,** Differential analysis (Δ) of the frequency of endocytic genes (EG; high-risk group; $n$ = 61) between patient clusters with high (C2, C4) versus low recurrence (C1). Coloured spots indicate endocytic genes with greater dysregulation; Green = ΔEG(C4-C1) ≥ 0.4; Pink = ΔEG(C2-C1) ≥ 0.4; Blue = ΔEG(C4-C1) & ΔEG(C2-C1) ≥ 0.4.

**G,** Same as **F** but differential analysis between patient subcluster C4a (highest recurrence) and clusters 1 and 3 (lowest recurrence). Blue spots = mean ΔEG(C4a-C1) & ΔEG(C4a-C3) ≥ 0.3 ($n$ = 40).

Data presented as mean ± S.E.M., one-way ANOVA followed by Dunnett's post hoc test (**C**), or Kruskal-Wallis test (**E**) (ns, not significant; *$P < 0.05$; **$P < 0.01$; ***$P < 0.001$).

**A** <u>THCA: BRAF-like RAI-treated (*n* = 137)</u>

| Gene Classifier | Cut-Off value | Sensitivity% | Specificity% | ROC AUC | ROC q-value | DFS q-value | HR | 95% CI |
|---|---|---|---|---|---|---|---|---|
| 10 | 8.756 | 71.43 | 81.03 | 0.8206 | <0.0001 | <0.0001 | 9.131 | 3.527, 23.639 |
| 20 | 25.22 | 90.48 | 89.66 | 0.9179 | <0.0001 | <0.0001 | 52.782 | 12.187, 228.605 |
| 30 | 27.62 | 85.71 | 94.83 | 0.9319 | <0.0001 | <0.0001 | 57.265 | 16.489, 198.873 |
| 40 | 37.82 | 95.24 | 81.03 | 0.9397 | <0.0001 | <0.0001 | 61.305 | 8.212, 457.659 |

| Classifier | Genes |
|---|---|
| 10 | *RAB35, SCAMP2, GRK6, CXCL16, AP2A2, NECAP1, CTTN, INPPL1, AP2A1, RAB5B* |
| 20 | *& CHMP7, CLTB, 1L10RA, HSPA8, ARRB2, GRN, DENND1A, SYNJ2, FCGR1A, SH3GL3* |
| 30 | *& FCGR3A, NECAP2, VAMP4, ARRB1, AGFG1, LILRB1, MYO6, OCIAD1, CYTH4, MYO1E* |
| 40 | *& TREM2, SIGLEC9, SIGLEC1, ARPC2, FCER1G, GRB2, SH3GL2, PIP5K1B, PIK3CA, SIGLEC7* |

**B**

| Gene Classifier | Validation datasets: BRAF MUT n = 228 (AUC, q-value) | RAI-treated n = 256 (AUC, q-value) | BRAF-like n = 260 (AUC, q-value) | THCA n = 486 (AUC, q-value) |
|---|---|---|---|---|
| 10 | 0.6845, 0.0023 | 0.6814, 0.0010 | 0.7229, 0.0003 | 0.5854, NS |
| 20 | 0.7669, < 0.0001 | 0.7366, < 0.0001 | 0.8092, < 0.0001 | 0.6336, 0.0030 |
| 30 | 0.7664, < 0.0001 | 0.7447, < 0.0001 | 0.8144, < 0.0001 | 0.6421, 0.0020 |
| 40 | 0.7675, < 0.0001 | 0.7508, < 0.0001 | 0.8116, < 0.0001 | 0.6373, 0.0025 |

**C**

| Patient Cohort | $N_{tot}$ | $N_{rec}$ | 30 Endocytic Gene Classifier: DFS *P*-value | DFS *q*-value | Univariate HR (95% CI) | Univariate *P*-value | 40 Endocytic Gene Classifier: DFS *P*-value | DFS *q*-value | Univariate HR (95% CI) | Univariate *P*-value |
|---|---|---|---|---|---|---|---|---|---|---|
| THCA | 486 | 46 | $4.14\times10^{-11}$ | $8.28\times10^{-11}$ | 5.717 (3.187-10.257) | $5.04\times10^{-9}$ | $8.99\times10^{-7}$ | $1.25\times10^{-6}$ | 3.861 (2.161-6.897) | $5.05\times10^{-6}$ |
| BRAF-like | 260 | 26 | $6.91\times10^{-17}$ | $3.11\times10^{-16}$ | 15.501 (6.604-36.383) | $3.05\times10^{-10}$ | $4.13\times10^{-10}$ | $7.43\times10^{-10}$ | 10.631 (4.241-26.650) | $4.63\times10^{-7}$ |
| BRAF-like, RAI-treated | 137 | 21 | $6.10\times10^{-28}$ | $1.10\times10^{-26}$ | 57.265 (16.489-198.873) | $1.86\times10^{-10}$ | $1.03\times10^{-13}$ | $3.71\times10^{-13}$ | 61.305 (8.212-457.659) | $5.99\times10^{-5}$ |
| RAI-treated | 256 | 34 | $2.01\times10^{-21}$ | $1.81\times10^{-20}$ | 13.680 (6.798-27.530) | $2.28\times10^{-13}$ | $3.86\times10^{-12}$ | $9.93\times10^{-12}$ | 8.421 (4.098-17.305) | $6.69\times10^{-9}$ |
| BRAF MUT | 228 | 27 | $5.51\times10^{-13}$ | $1.65\times10^{-12}$ | 10.953 (4.871-24.628) | $7.06\times10^{-9}$ | $4.39\times10^{-8}$ | $6.59\times10^{-8}$ | 7.767 (3.270-18.452) | $3.42\times10^{-6}$ |
| BRAF MUT, RAI-treated | 125 | 21 | $2.48\times10^{-20}$ | $1.49\times10^{-19}$ | 38.250 (11.139-131.341) | $7.06\times10^{-9}$ | $8.75\times10^{-12}$ | $1.97\times10^{-11}$ | 51.72 (6.930-385.975) | $1.19\times10^{-4}$ |
| RAS-like | 115 | 8 | 0.504 | 0.504 | NS | 0.661 | 0.376 | 0.398 | NS | 0.565 |
| Non RAI-treated | 167 | 6 | 0.262 | 0.295 | NS | 0.480 | 0.146 | 0.175 | NS | 0.379 |

**D** 30 Endocytic Gene Classifier

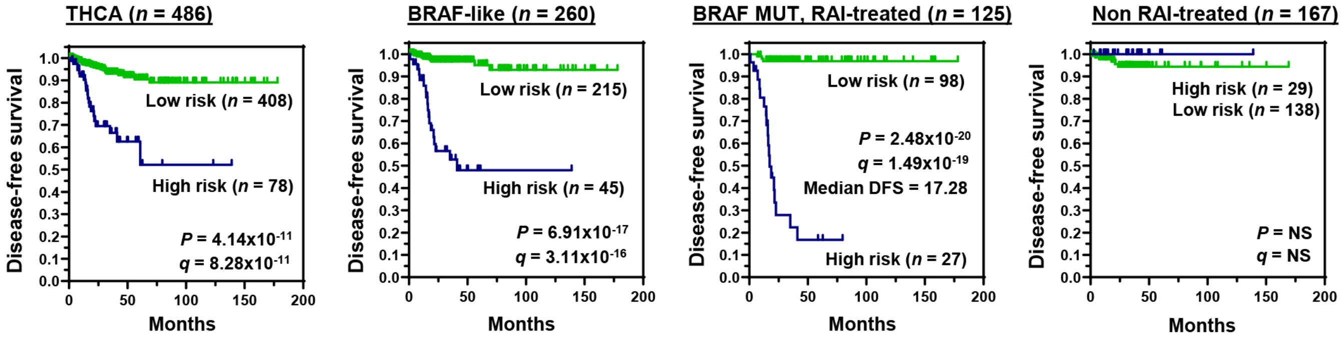

**Figure S7.** Construction and validation of endocytic multigene classifiers for PTC recurrence.

**A,** Prognostic profile of endocytic risk score multigene classifiers (*n* = 10 – 40 genes). Estimations of sensitivity and specificity (%) for the BRAF-like, RAI-treated THCA cohort (*n* = 137) are given, as well as cut-off expression values for stratification. AUC, area under curve; DFS, disease-free survival; HR, hazard ratio; CI, confidence interval. (*lower*) Genes included in the various classifiers are indicated. Gene order based on consolidation of the 40 gene risk score, in which genes were ranked on the relative proportion of their contribution towards the overall risk score value.

**B,** ROC analysis (AUC) of different gene classifiers (*n* = 10 - 40) in validation larger THCA datasets as indicated. Red box indicates highest AUC value for each THCA cohort.

**C,** Comparison of Kaplan-Meier and univariate Cox regression analysis in different THCA cohorts using either a 30- or 40 endocytic gene risk score classifier. Total number of patients ($N_{tot}$) and recurrent cases ($N_{rec}$) in each patient cohort are given.

**D,** Representative Kaplan-Meier analysis of DFS for the THCA (*n* = 486), BRAF-like (*n* = 260), BRAF MUT RAI-treated (*n* = 125) and Non RAI-treated (*n* = 167) THCA cohorts stratified using the 30 endocytic gene classifier; log-rank test. Number (n) of patients per expression sub-group (high/low), *P*-values and *q*-values are shown.

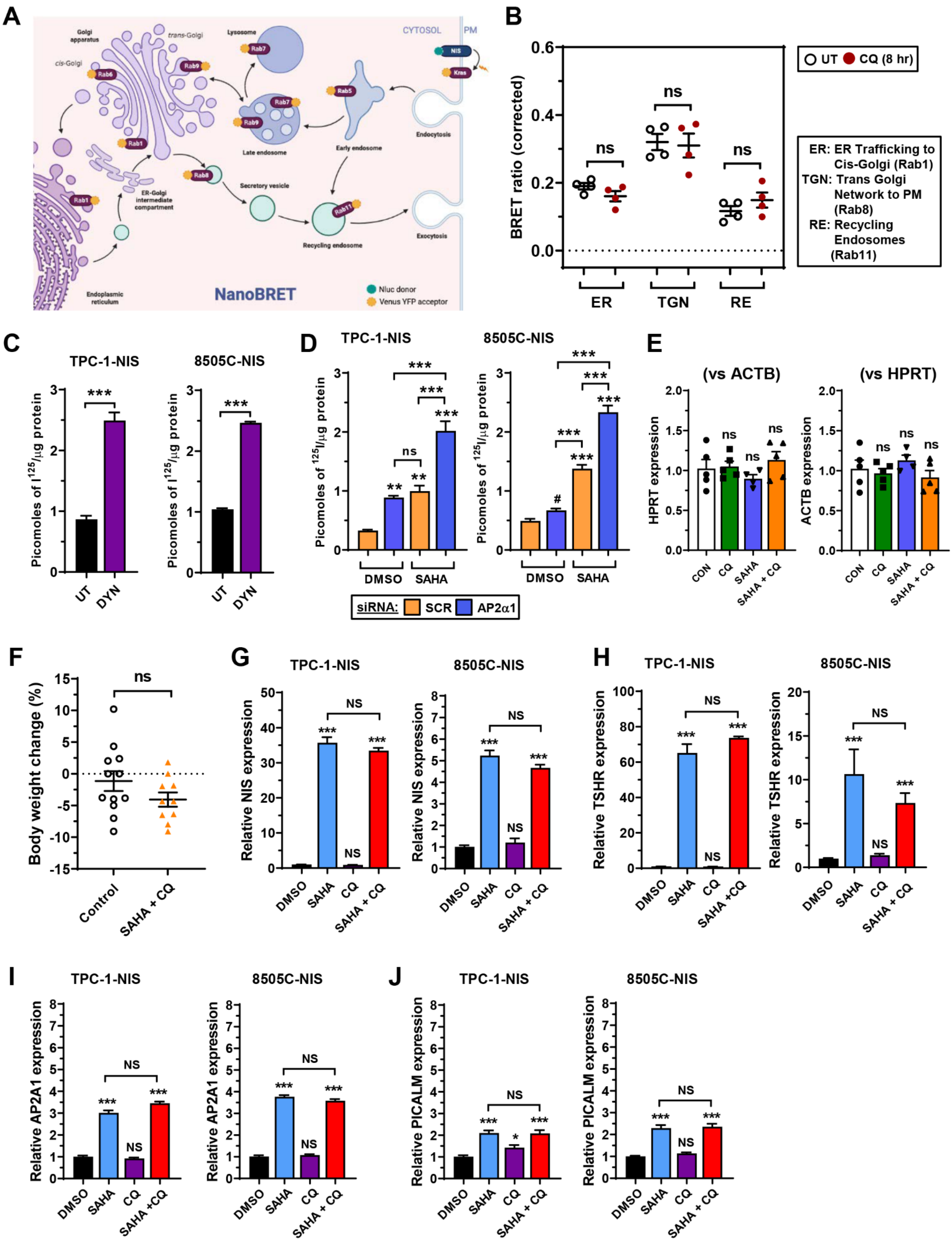
A
Golgi apparatus
trans-Golgi
cis-Golgi
Lysosome
CYTOSOL
PM
NIS
Kras
Rab6
Rab9
Rab7
Rab5
Rab1
Rab8
Rab11
Late endosome
Early endosome
Endocytosis
ER-Golgi intermediate compartment
Secretory vesicle
Recycling endosome
Exocytosis
Endoplasmic reticulum
NanoBRET
Nluc donor
Venus YFP acceptor
B
BRET ratio (corrected)
UT
CQ (8 hr)
ER
TGN
RE
ns
ER: ER Trafficking to Cis-Golgi (Rab1)
TGN: Trans Golgi Network to PM (Rab8)
RE: Recycling Endosomes (Rab11)
C
TPC-1-NIS
8505C-NIS
Picomoles of I125/μg protein
UT
DYN
D
DMSO
SAHA
siRNA: SCR AP2α1
E
(vs ACTB)
(vs HPRT)
HPRT expression
ACTB expression
CON
CQ
SAHA
SAHA + CQ
F
Body weight change (%)
Control
SAHA + CQ
G
Relative NIS expression
DMSO
NS
H
Relative TSHR expression
I
Relative AP2A1 expression
J
Relative PICALM expression

**Figure S8.** Transcriptional drug responses in thyroid cancer cells.

**A,** Schematic depicting NanoBRET assay to monitor close proximity of NIS with different subcellular markers (e.g. Rab 1, 8 and 11) tagged with Venus. Created with BioRender.com.

**B,** Profiling subcellular changes of NIS using the NanoBRET assay in CQ-treated HeLa cells. HeLa cells were transiently transfected with NIS tagged with NLuc, and one of the subcellular markers Rab1 (ER trafficking to cis-golgi), Rab8 (Trans golgi network to PM) or Rab11 (Recycling endosomes) tagged with Venus.

**C,** RAI uptake in TPC-1-NIS and 8505C-NIS cells following Dynasore (DYN) treatment.

**D,** RAI uptake in TPC-1-NIS and 8505C-NIS cells following AP2α1-siRNA depletion and SAHA treatment. Scr – scrambled control siRNA.

**E,** HPRT and ACTB mRNA levels in thyroid glands dissected from WT BALB/c mice administered with CQ and SAHA either alone or in combination.

**F,** Body weight change (%) in WT BALB/c mice administered with a combination of CQ and SAHA ($n$ = 10) versus controls ($n$ = 12).

**G-J,** Relative NIS (**G**), TSHR (**H**), AP2A1 (**I**) and PICALM (**J**) mRNA levels in TPC-1-NIS and 8505C-NIS cells treated with CQ and SAHA either alone or in combination.

Data presented as mean ± S.E.M., $n$ = 3-7 unless stated, one-way ANOVA followed by Tukey's post hoc test (ns, not significant; $^{*}P < 0.05$; $^{**}P < 0.01$; $^{***}P < 0.001$) or unpaired two-tailed t-test ($^{\#}P < 0.05$).
